# Disease associated mutations in tau encode for changes in aggregate structure conformation

**DOI:** 10.1101/2023.04.24.537889

**Authors:** Kerry Sun, Tark Patel, Sang-Gyun Kang, Allan Yarahmady, O. Julien, Jonathan Heras, Sue-Ann Mok

## Abstract

The accumulation of tau aggregates is associated with neurodegenerative diseases collectively known as tauopathies. Tau aggregates isolated from different tauopathies such as Alzheimer’s disease, corticobasal degeneration and progressive supranuclear palsy have distinct cryo-electron microscopy structures with respect to their packed fibril cores. To understand the mechanisms by which tau can be sensitized to form distinct aggregate conformations, we created a panel of tau variants encoding for individual disease-associated missense mutations in full-length 0N4R tau (wild-type and 36 mutants). We developed a high-throughput protein purification platform for direct comparison of tau variants in biochemical assays. Structural analysis of the protease-resistant core of tau aggregates formed *in vitro* reveals that mutations can promote aggregate core packing distinct from that produced by WT tau. Comparing aggregate structure changes with aggregation kinetic parameters for tau mutants revealed no clear linkage between these two aggregation properties. We also found that tau mutation-dependent alterations of tau aggregate structure are not readily explained by current tau fibril structure data. This is the first study to show the broad potential of tau mutations to alter the packed core structures contained within aggregated tau and sheds new insights into the molecular mechanisms underlying the formation of tau aggregate structures that may drive their associated pathology in disease.

## Introduction

Tauopathies are a group of neurodegenerative disorders that feature the pathological accumulation of aggregates containing microtubule-associated protein tau^1,2^. The fibrillar forms of tau aggregates are characteristically composed of a core region containing regularly tightly packed monomer sequences rich in beta-sheet structure that are surrounded by more loosely packed or disordered N and C-terminal sequences termed the “fuzzy coat”^3–5^. Protease digestion of tau fibrils isolated from patient samples defined the sequences protected within the tightly packed core and suggested that 1) tau can adopt multiple conformations in fibril structures and 2) specific fibril structures tracked with individual tauopathies: Alzheimer’s disease (AD), Pick’s disease (PD), corticobasal degeneration (CBD), and progressive supranuclear palsy (PSP)^6–15^. The distinct features of tau fibril core structures from multiple tauopathies have now been confirmed to atomic resolution by Cryo-electron microscopy (Cryo-EM)^10–15^. Prion-like properties have been ascribed to tau^16,17^ in which the conformation of the tau aggregate promotes aggregate propagation via cell-to-cell spreading and faithful recruitment and templating of its structure to native tau monomers^18–21^. Tau aggregates isolated from tauopathies and introduced to cells display differential requirements for templating aggregation^22^ and AD, CBD, or PSP fibrils are injected into mice they promote distinct patterns of pathology that share qualities with the corresponding human disease^23^. Thus, understanding the mechanisms by which distinct tau aggregate structures are generated and how they are linked to tauopathy would contribute to a better molecular understanding of the genesis and progression of these diseases.

Over 50 mutations in the *MAPT* gene that encodes tau have been linked to tauopathies^24^. The collection of mutations show heterogeneity with respect to penetrance, age of onset, clinical phenotypes, and their association with individual tauopathies^24–29^. At the molecular level, individual tau mutations have also been shown to differentially modulate tau biochemical properties, cellular dysfunction associated with pathogenesis. For example, intronic mutations (and some missense mutations) alter the regulation of tau splicing and the resulting composition of tau isoforms expressed^30–32^. Individual missense mutations have been shown to differ from wild-type (WT) in one or more properties including aggregation propensity^33–35^, cellular aggregate seeding^36,37^, microtubule binding and dynamics^35,38–41^, protein interaction partners^42,43^, post-translational modifications^44,45^, engagement and processing by the protein quality control machinery^43,46–48^ and axonal functions^49,50^. Mice expressing individual tau missense mutants produce distinct phenotypic profiles in regards to cellular dysfunction, patterns and timing of tau aggregate pathology, and behavioral deficits^51^. In light of the evidence of multiple tau aggregate conformations linked to tauopathies, we reasoned that one a potential factor that could contribute to the pathogenic effects of tau mutations is their ability to promote the formation of alternate tau aggregates that have not been extensively investigated.

Although there have been previous studies of structural differences induced by a small subset of mutations in short synthetic constructs of tau^52,53^, comparison of the effects of a broad set of tau mutations in the context of full-length tau has not been characterized. Thus, to investigate the role of disease-associated tau missense mutations in modulating tau aggregate structure, we generated a panel of 37 tau variants (WT and 36 missense mutants) for systematic comparison of their properties. We first developed a high-throughput approach for expression and plate-based purification of recombinant tau mutant proteins. We then used trypsin digestion assays to profile the protected core regions of individual tau mutant aggregates generated *in vitro*. Our high-throughput system also enabled us to perform cross comparison of our aggregates structure profiles with other aggregation properties such as aggregation kinetics. We found that multiple disease-associated mutations formed different aggregate structures than WT tau which could explain their ability to cause pathogenesis despite having WT-like aggregation kinetics. Our dataset also showed that current structural data on tau fibrils did not effectively predict the mutation-dependent changes in aggregate structure that we observed. This study is a proof of concept that our experimental platform can be used and adapted to systematically tease out the complex relationships between tau sequence, aggregation and cellular dysfunction.

## Materials and Methods

### Cloning of tau constructs

Mimics of disease-associated tau mutations were generated in the 0N4R isoform of human tau by site-directed mutagenesis. The parent vector is pET28 0N4R which encodes for expression of WT 0N4R tau that is N-terminally fused to a sequence containing a 6XHis-tag and TEV cleavage site. Q5 polymerase (NEB) was used to amplify the pET28 0N4R plasmid^46^ with desired mutagenesis primer pairs. Amplified templates were treated with DpnI (ThermoFisher) prior to transformation into DH5ɑ competent cells (Agilent). Successful mutation of isolated clones was verified by Sanger sequencing and clones were retransformed into BL21-CodonPlus (DE3)-RP competent cells (Agilent) for protein expression.

### Tau purification

Large-scale protein purifications of tau were carried out as previously described in Mok et al^46^. The large-scale purification protocol was modified as followed for small-scale protein expression and purification. Cells transformed with individual tau variants were grown to an O.D. 600 of 0.7 then induced with IPTG (500 µM) for 3hr at 30°C to allow for protein expression. Pelleted cells arrayed in 96-well plates were resuspended in resuspension buffer (1.5 mM KH_2_PO_4_, 8 mM Na_2_HPO_4_, 2.7 mM KCl, 2 mM DTT, 2 mM PMSF, pH 7.2) then lysed with lysozyme (0.3 mg/ml final) in 3 successive freeze-thaw cycles. Lysates were treated with 12 U/mL of Benzonase for 30 minutes at 4°C then boiled for 20 minutes and cleared by 2 rounds of centrifugation at 3214xg for 15 minutes. Tau was purified from the supernatant fraction by in-well chromatography using SP ImpRes/SP sepharose FF resin (Cytiva) incubated in 96-well filter plates (Supor PES membrane 1.2 µM, Pall). Following 4 rounds of resin exchange in wash buffer (1.5 mM KH_2_PO_4_, 8 mM Na_2_HPO_4_, 2.7 mM KCl, 2 mM DTT, 50-150 mM NaCl, pH 7.2), tau was eluted in (1.5 mM KH_2_PO_4_, 8 mM Na_2_HPO_4_, 2.7 mM KCl, 2 mM DTT, 350 mM NaCl, pH 7.2). Eluted tau protein was subject to buffer exchange in low salt buffer (1.5 mM KH_2_PO_4_, 8 mM Na_2_HPO_4_, 2.7 mM KCl, 2 mM DTT, pH 7.2) then aggregation assay buffer (Dulbecco’s PBS, pH 7.4, 2 mM DTT) using 3K MWCO filter plates (Pall). Buffer exchange was monitored by measuring conductivity of the protein sample with a conductivity probe calibrated to standard solutions (INLAB 751-4MM, Mettler Toledo). Concentrations of purified tau protein were determined using a reducing agent compatible BCA assay (Thermo Fisher). Protein purity of final preparations was assessed by coomassie-stained SDS-PAGE or MALDI-TOF Mass spectrometry (Bruker Autoflex Speed MALDI-ToF, Bruker Daltonic).

### Tau kinetic aggregation assays

Aggregation kinetic assays were performed and analyzed as previously described^46^. Briefly, reactions consisted of 10 µM tau, 10 µM thioflavin T (Sigma), 44 µg/mL heparin sodium salt (Santa Cruz) in assay buffer. Aggregation reactions were carried out at 37 °C with continuous shaking and monitored via ThT fluorescence (excitation, 444 nm; emission, 485 nm; cutoff, 480 nm) in a Spectramax M5 microplate reader (Molecular Devices). Readings were taken every 5 min for a minimum of 24 h. For analysis of kinetic aggregation curves, individual baseline subtracted curves were fitted to the Gompertz function^54,55^ to extract kinetic parameters (lag time, apparent elongation rate constant, amplitude). A weighting of 1/Y was applied during the fitting to accurately capture lag time values.

### Trypsin digestion of tau aggregates

Tau aggregates directly sampled from *in vitro* aggregation assays (15 µL) were treated with a final concentration of 0.03 mg/mL mass spectrometry-grade trypsin (ThermoFisher) in D-PBS. A time course digestion using WT 0N4R tau as the substrate was performed for each trypsin lot to calibrate assay conditions (total digestion time) across experiments. Trypsin digest assays were incubated at 37°C with shaking at 800 rpm for the indicated times. Digestion reactions were stopped by addition of sample buffer (Fluorescent Master Mix, ProteinSimple) and heating to 95°C for 5 min. Tau fragments were resolved using the Jess capillary gel electrophoresis system (ProteinSimple) with the 2-40 kDa separation. Reagents and equipment were purchased from ProteinSimple unless stated otherwise. Four microliters of each sample were loaded into the top-row wells of plates preloaded with proprietary electrophoresis buffers designed to separate proteins of 2-40 kDa. Subsequent rows of the plate were filled with blocking buffer, primary and secondary antibody solutions, and chemiluminescence reagents, according to the manufacturer’s instructions. Detection of protein fragments was performed with the total protein detection module or capillary western blots using anti-tau antibodies^56^. Primary antibodies were anti-tau monoclonal antibodies including DA9 (aa 102–140, 1:10 dilution), ET3 (aa 273-288, 1:10 dilution)^57^ and 77G7 (BioLegend, aa 316-355, 1:100 dilution). DA9 and ET3 antibodies were generously provided by Peter Davies. Secondary antibodies were anti-mouse secondary HRP use according to manufacturer’s directions (ProteinSimple). CompassSW software (ProteinSimple) was used to generate chromatograms representing the lane profiles of separated protease-resistant tau fragments detected using the total protein module (ProteinSimple) or tau antibodies. Chromatograms plot signal intensity versus apparent MW, calibrated using protein standards included in each capillary run.

### Comparative analysis of trypsin-resistant tau fragment profiles

To perform a cross-comparison of chromatograms (fragment banding patterns) a normalization step was first conducted to generate standardized increment values with respect to kDa across all chromatograms. Namely, from each profile, a 1-D function is interpolated, and values in the range from 6 to 24 with a step of 0.1 are obtained, see Supp. Fig. 1.

Subsequently, the distance among the interpolated profiles is obtained by computing the correlation distance. Given two 1-D arrays, *u* and *v*, the correlation distance between *u* and *v*, is defined as

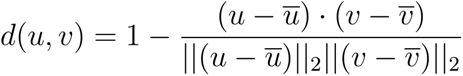

where ū is the mean value of the elements of *u*, and *x*·*y* is the dot product of *x* and *y*. Once that the distance among the profiles is computed, they are grouped by hierarchical clustering using average linkage (this is represented by means of a dendrogram)^58^, and the distance among the samples is graphically shown by using a heatmap. This process has been implemented in the Python programming language and using the scikit-learn^59^ and scipy^60^ libraries.

### Epitope mapping of trypsin resistant tau fragments

A script was generated in python programming language to identify potential tau sequences corresponding to individual trypsin-resistant tau fragments based on their reactivity to assayed tau antibodies and their apparent MW via capillary gel electrophoresis. The script identifies potential tau fragment sequences generated by trypsin digestion for a given MW. It then includes/excludes potential sequences based on their reactivity with assayed tau antibodies. The program also searches for potential tau dimer sequences if no matching sequence is predicted from the initial analysis.

### Dot blots

0N4R and 0N3R WT tau were expressed and purified in alternating wells of a 96-well plate by small-scale tau expression and purification. 100 pmol of purified tau sample from each well were applied directly to a 0.1μm nitrocellulose blotting membrane. The membrane was probed with primary antibodies including anti-tau mouse ET3 (1:1000 dilution) and 6X - His Tag rabbit monoclonal antibody (Invitrogen, 1:3500 dilution). The secondary antibodies used were: Cytiva CyDye 700 goat-anti-mouse and Cytiva CyDye 800 goat-anti-rabbit (diluted 1:10000). Following secondary antibody incubation, the blot was visualized with a LICOR imager using the 700 nm and 800 nm channels.

### MALDI-TOF

For MALDI analysis 1μL of each sample was mixed with 1μL of sinapinic acid (10 mg/ml in 50% acetonitrile/water + 0.1% trifluoroacetic acid). 1μL of the sample/matrix solution was then spotted onto a stainless steel target plate and allowed to air dry. Mass spectra were obtained using a Bruker Autoflex Speed MALDI-TOF (Bruker Daltonic GmbH). All MS spectra were recorded in positive linear mode and external calibration was performed by use of a standard protein mixture. Predicted m/z for full-length protein = 43801.1696 (with met oxidation= 43961.1188).

### Modeling tau mutations into existing Cryo-EM tau fibril structures

PDB structures and their associated 2FoFc maps were modelled in PyMOL (AD SF: 5O3T, AD PHF: 6HRE, CBD: 6TJO, PSP: 7U0Z, snake: 6QJH, twister: 6QJM, jagged: 6QJP). Mutations were modelled onto structures using the mutagenesis function and all backbone-dependent rotamers were analyzed. Amino acid mutations where all allowed side chain rotamers clash with the existing structure were deemed as incompatible.

### Code availability

All code described in Materials and methods for processing data from aggregation kinetics, digest profile comparisons and epitope mapping are available at the following GitHub repository: https://github.com/sueannmok/tau_digest_and_kinetics_tools.git

## Results

### Validate a high-throughput tau purification platform

Our first objective was to be able to directly compare our 37 disease-associated tau mutants with respect to multiple aggregation properties. Tau transcripts are subject to alternative splicing leading to the production of multiple tau isoforms. Six major tau isoforms are prevalent in the adult CNS and differ in the inclusion/exclusion of two N-terminal inserts (0N, 1N or 2N) and three versus four microtubule binding repeats (3R, 4R). For our study, we targeted introduction of familial mutations into the 0N4R tau isoform because it is a major component of tau aggregates in multiple tauopathies (AD, PSP, CBD)^61,62^ and it aggregates relatively rapidly *in vitro*^46^. We developed a platform for purifying recombinant human 0N4R tau protein in a 96-well plate format (Fig. 1A) so that all individual tau mutants could be purified in parallel and the entire mutant panel could be assayed across multiple protein preparations. Our small-scale tau purification protocol retained the major procedures reported in traditional large-scale methods including protein expression in *E.coli*, boiling of the lysate, cation exchange and buffer exchange into standard conditions (DPBS, 2mM DTT) suitable for protein aggregation assays^33,46,63^. SDS-PAGE of small-scale purified WT 0N4R tau was the expected full-length form, similar to the same tau construct obtained from a traditional large-scale purification protocol (Fig. 1B). MALDI-TOF analysis of small-scale purified 0N4R WT tau detected a single major species at 44054 m/z closely corresponding to the predicted full-length product (Fig. 1C). To test the potential for cross-contamination of proteins between individual wells, we purified two different isoforms of tau (0N3R and 0N4R) arranged in alternating wells within the plate (Fig. 1D). The purified samples were subjected to a dot blot with an antibody specifically recognizing the 0N4R isoform. Antibody reactivity was only detected in the purified 0N4R samples and not the 0N3R sample purified from adjacent wells suggesting minimal/no cross-contamination occurs between wells during purification. Each in-well purification yielded on average >200 μg of purified tau prepped for downstream assays (Supp. Fig. 2A).

**Figure 1.**
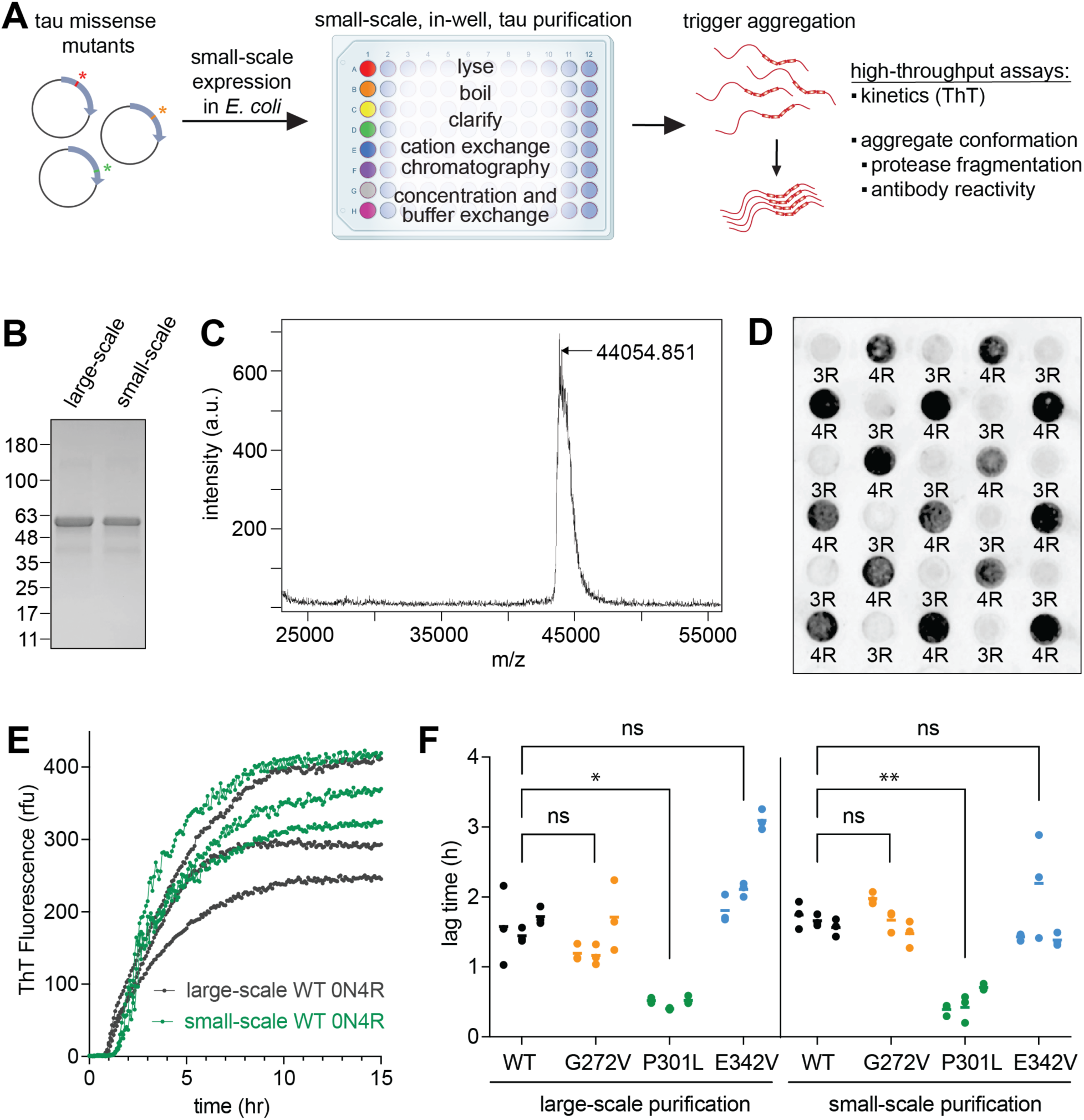
Validation of high-throughput, small-scale purification and aggregation platform for recombinant tau. A) Schematic outlining major components of the tau purification and downstream assay platform. Individual tau constructs with encoded disease-associated mutations are expressed in small volume *E. coli* cultures. Each culture is transferred to individual wells of a series of 96-well plates during the lysis and purification procedure to obtain purified tau. The purified tau is in D-PBS with 2mM DTT compatible with all downstream assays used in this study. B) SDS-PAGE comparison of purified WT 0N4R tau using a traditional large-scale protocol or our small-optimized scale protocol. One μg of tau is loaded per lane and separated proteins are detected by Coomassie blue staining. The calculated MW of the expressed tau is 44 kDa however, the protein migrates at an apparent MW of ~50 kDa due to a high number of positively charged residues contained within the sequence. Results are representative of 3 individual protein preparations. C) Mass spectrum of small-scale purified tau demonstrating a single peak corresponding to the calculated mass of the full-length protein. Results are representative of 3 individual protein preparations. D) Dot blot demonstrating no cross-contamination of tau samples during small-scale purification. Cultures expressing 0N4R or 0N3R WT tau were purified in alternating wells of the 96-well purification platform. Each purified tau sample was transferred to a dot blot while maintaining its plate position relative to other samples during purification. Dots corresponding to 0N4R and 0N3R tau samples are labelled as 4R and 3R, respectively. The dot blot was probed with a 4R tau antibody (ET3) that recognizes an epitope found in 4R but not 3R tau isoforms. A replicate blot probed with a His-tag antibody was used to verify loading of purified tau as shown in Supp Fig. 1D. E) Aggregation kinetic curves obtained for WT 0N4R tau purified using large-scale versus small-scale methods. Aggregation reactions were initiated by introducing heparin to 10 μM tau and the reaction was monitored with ThT. The individual data points for reactions from three technical replicates are plotted. F) Comparison of calculated lag time values for WT 0N4R tau and mutants (G272V, P301L, S342V) purified via large or small-scale protocols. For each tau variant, individual lag time values for three technical replicates (aligned dots), assayed from three individual protein preparations (side-by-side columns of dots) are plotted along with the mean ± SD for each experiment (bar). *p = < 0.05, **p = <0.01, nested one-way ANOVA, post-hoc Dunnett’s test.

Individual disease-associated mutations were introduced into human 0N4R tau by site-directed mutagenesis and purified using our small-scale purification platform. We first compared large-scale and small-scale purified tau in an *in vitro* aggregation assay for a subset of tau mutants (WT, G272V, P301L, E342V). The aggregation kinetic assays were carried out in a miniaturized 384-well format that we previously developed^46^ and require less than 9 μg of tau protein per reaction. Tau aggregation was tracked with the fluorescent amyloid dye Thioflavin T (ThT) following the addition of a common aggregation accelerant, heparin. An example of the kinetic aggregation curves obtained for large-scale versus small-scale purified WT 0N4R tau is shown in Fig. 1E. Individual aggregation curves were fitted to the Gompertz function to extract kinetic parameters: lag time, elongation rate constant, amplitude. Each tau mutant was assayed across three different protein preparations for each purification method. Between tau mutants tested, similar relationships were observed with respect to the extracted kinetic parameters (Fig. 1F, Supp. Fig. 2B-C). For example, G272V tau consistently aggregated with a lag time similar to WT whereas P301L tau had a significantly shorter lag time than WT (Fig. 1F). Moreover, calculated kinetic parameters showed similar variability when comparing large-scale versus small-scale purified tau aggregated from different preparations. Overall, these results support that tau purified using small-scale methods is comparable to tau purified by classical large-scale methods and critically, allows us to perform cross-comparisons of tau mutants for their aggregation properties, at a higher throughput capacity.

### Tau disease-associated mutations drive the formation of aggregate structures distinct from WT

We next determined if single tau mutations were sufficient to promote the formation of alternative aggregate structures. As a readout of aggregate structure, we used protease digestion of aggregated tau to identify tau fragments buried in the tightly packed aggregate core which are more protected from protease activity. The specific pattern of protected tau fragments has been used to detect distinct aggregate species in both recombinant tau and disease patient samples^6,7,21^. We first explored the kinetics of digestion for WT 0N4R tau with the protease, trypsin. The tau protein contains trypsin sites spread throughout its sequence at an average of one site every 8 residues making it a robust sensor of changes in tau sequence access/structure (Supp. Fig. 3A). WT 0N4R tau was incubated with heparin and allowed to aggregate for 24 h prior to direct digestion of the reaction with mass spectrometry grade trypsin (has enhanced stability and no chymotrypsin activity) for 0, 1, 2, 3, 6, and 18 h. Capillary gel electrophoresis technique combined with total protein detection was used to quantify the migration and relative amounts of trypsin-resistant fragments. The technique is highly sensitive and can detect the fragments produced from as little as 0.8 μg of digested tau aggregate sample. Fig 2A displays the simulated lane view of the fragments for individual samples and Fig. 2B shows the chromatogram profiles of the corresponding samples. Serendipitously, trypsin itself is not detected in our profiles since the mass spectrometry grade trypsin is modified at lysine residues utilized by the total protein detection method. Monomer tau treated with trypsin was completely digested to 2-3 kDa fragments within 1 h trypsin treatment. Undigested tau aggregates migrated at >40 kDa corresponding to the expected size of full-length tau. Loss of full-length tau after trypsin addition coincided with the generation of smaller peptide fragments. At 1 h and 2 h digest timepoints, the pattern of produced fragments was fairly consistent giving three major fragment peaks at 10, 16, and 28 kDa and confirmed the presence of a stable core that is relatively refractory to trypsin digestion of its internal sequences. Similar to the digested monomer tau control sample, a major peak at 2 kDa was also observed. Since our aggregated samples were not further purified prior to trypsin treatment, the 2 kDa band likely corresponds to unaggregated tau present within our original reactions. At 3, 6, and 18 h time points, a shift of 1-2 kDa towards smaller MWs was observed for each of the major fragments. For example, at the 3 h digest timepoint, a 9 kDa band was detected along with the 10 kDa band. By 6 h, only the 9 kDa band was the major peak. These results likely suggest prolonged trypsin treatment leads to increased digestion from the core fragment ends. Based on these results, digestion reactions were kept to <2 h for the remainder of the study.

**Figure 2.**
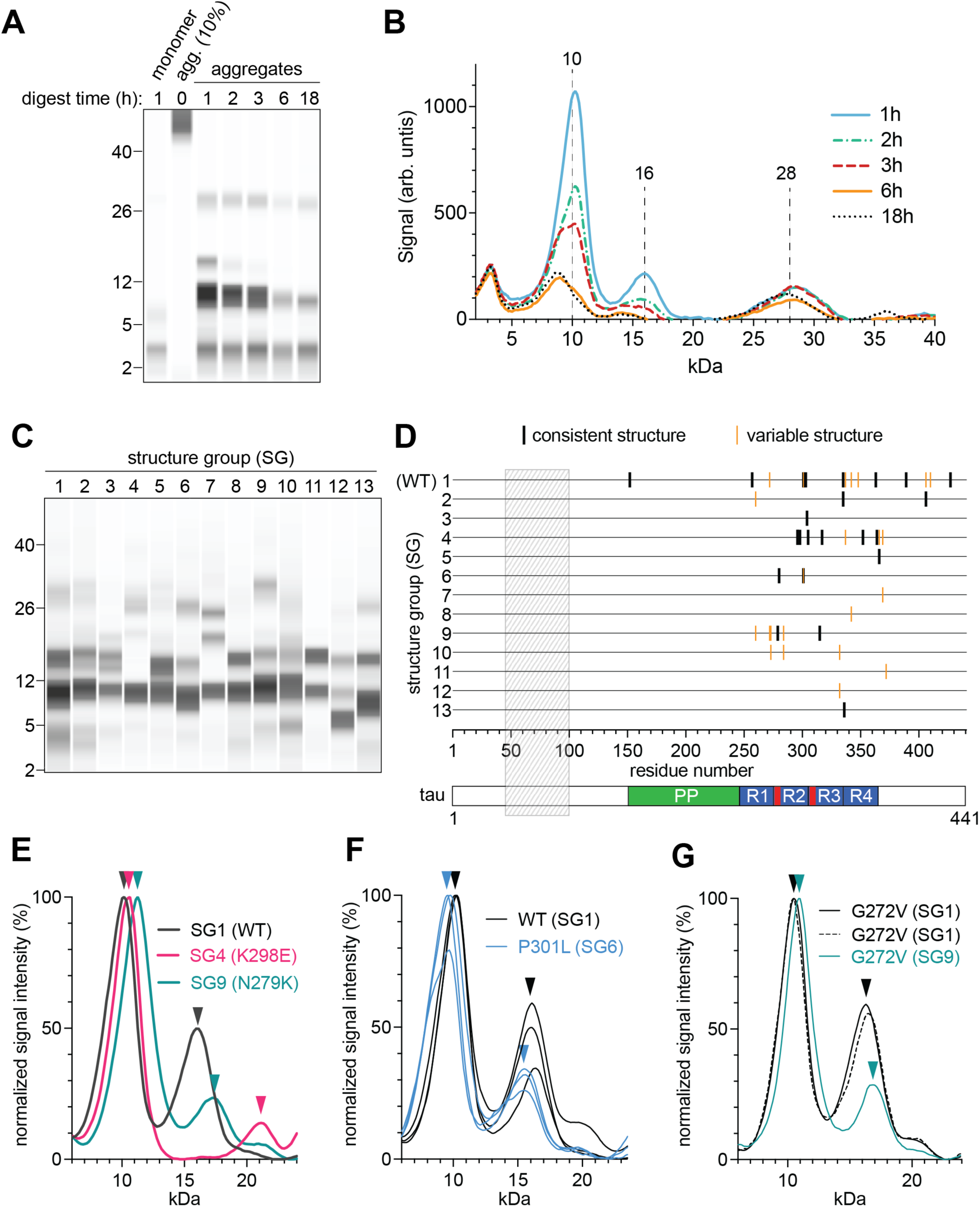
Tau mutations promote the formation of tau aggregate structures distinct from WT as indicated by altered trypsin digest fragment profiles. A-B) Time course of trypsin digestion of WT 0N4R tau aggregates. WT 0N4R tau aggregates were incubated with trypsin for the indicated time or left untreated (0 h). A control sample of tau monomer treated with trypsin for 1 h was also prepared. Samples were processed for capillary gel electrophoresis and total protein detection (lysine-dependent labeling) of protease-resistant fragments. Note that trypsin is not detected due to lysine modification of the enzyme. Results are representative of three independent experiments. A) Capillary lane view of separated trypsin-resistant fragments. Migration of MW standards are indicated. B) Corresponding chromatograms for samples digested with trypsin for 1, 2, 3, 6, and 18 h from A). Major peak maxima within spectra indicated by vertical dashed lines and corresponding MW. C) Representative capillary lane view of digest profiles for each structure group identified (1-13). Migration of MW standards are indicated. D) 0N4R tau mutants plotted according to their location within the tau sequence (x-axis) and structure group classification (y-axis). The numbering on the x-axis is based on the longest isoform of tau 2N4R. A hatched box covers the sequence region not included in the 0N4R tau isoform constructs used in this study. Tau mutants that gave a single digest profile across three trials were termed as having a consistent structure (black line) whereas those that gave an alternate profile in at least one experiment were classified as having variable structure (orange lines, all structure groups indicated for each mutant). A schematic of the tau sequence is included below the graph. The location of major sequence regions are indicated: PP = polyproline region, R1-R4 = microtubule binding repeat regions, red boxes = aggregation motif locations. E-G) Chromatograms of digested fragment profiles over the 6-24 kDa region. For the y-axis, signal intensity is normalized with the maximal signal value set to 100%. The structure group and corresponding tau mutant producing the fragment profile plotted are indicated by color. Color coordinated arrowheads mark the maxima of major peaks for each fragment profile. E) Representative chromatograms of three major structure groups defined in this study: SG1, SG4, and SG9. F) Individual profiles produced from independent experiments (n=3) for WT 0N4R tau and P301L 0N4R tau. G) Individual profiles for the variable structure mutant, G272V 0N4R tau.

Using our digest conditions, we directly compared the protease-resistant fragment profiles for aggregates generated from WT tau and 36 disease-associated tau mutants. Tau variants were small-scale purified in parallel, aggregated and trypsin digested with the same reaction conditions. We observed that a subset of tau mutants produced a digest fragment profile that differed from WT tau. To systematically compare the trypsin digest fragment profiles of individual samples and identify distinct profiles, we developed an algorithm to process gel chromatogram data and perform hierarchical cluster analysis (see Materials and Methods for details). Clustering is based on similarity of the overall shape of the chromatogram profile (relative peak heights and distances). We restricted analysis to the 6-24 kDa range which we observed more consistently detected signals from fragments above background levels. We applied the analysis to a dataset containing three independent digest reactions for each tau variant within our panel. Using a 90% of threshold for similarity, the dataset was binned into 13 separate profiles (Supp. Fig. 4) that we term “structure groups” (SG). The representative fragment profile for each structure group is presented in Fig. 2C. The profile for WT tau corresponds to the pattern for structure group 1 (SG1) and the remaining structure groups are ordered as a function of decreasing similarity score relative to SG1. Fig. 2D shows a schematic of the distribution of tau mutants in each structure group and their relative location within the tau sequence. In the corresponding table (Table 1), each tau mutant its identified structure group(s) is indicated.

**Table 1.**
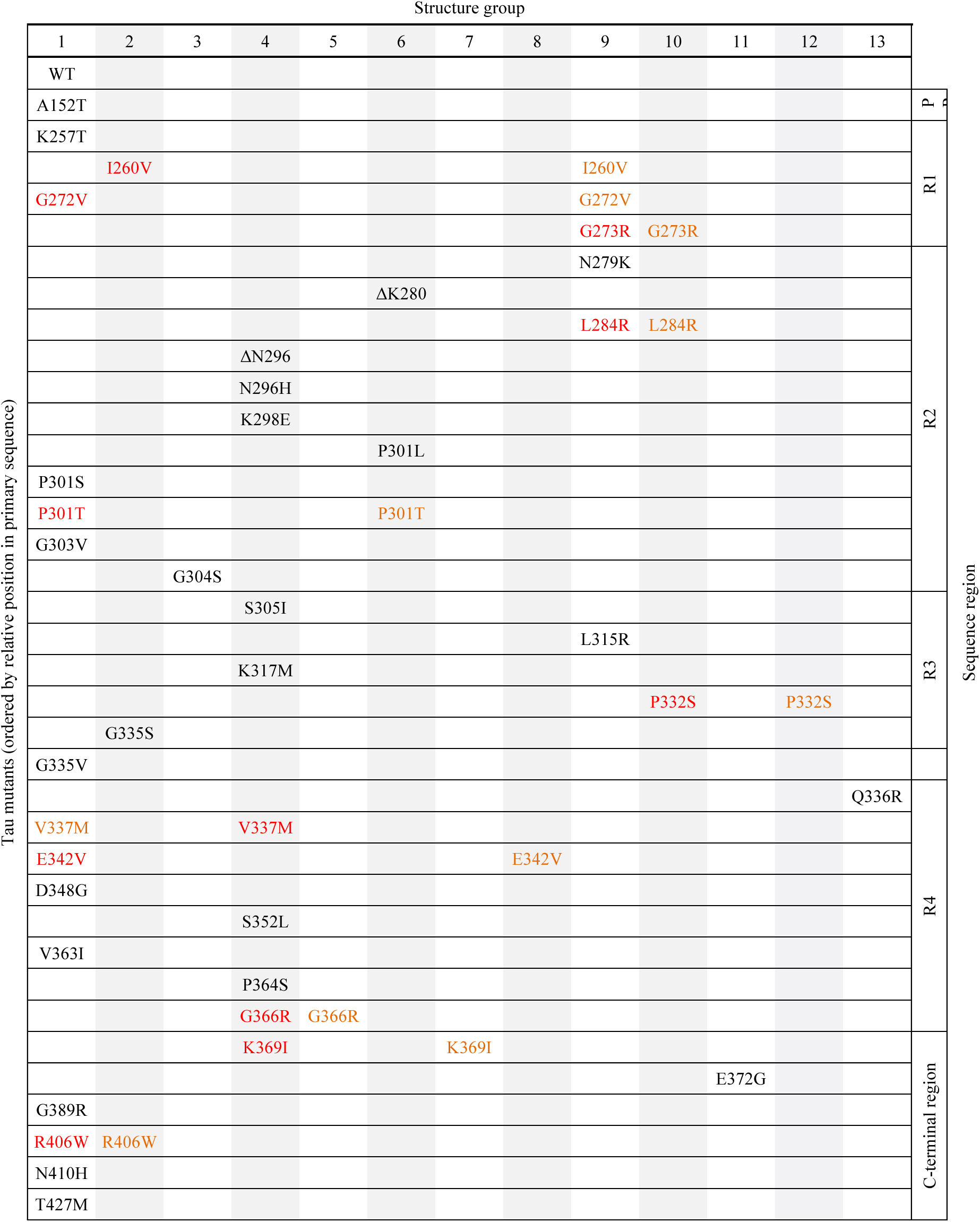
Structure group classifications of tau mutants based on their trypsin-resistant fragment profiles. Each tau mutant is binned into the indicated structure groups (1-13). Mutants are ordered from top to bottom based on their relative sequence location. Mutants that gave a consistent structure group across all three independent experiments are colored in black. Mutants that binned into more than one structure group are colored red (occurred in 2 experiments) and orange (occurred in 1 experiment). The tau sequence region associated with each mutant is indicated at the far right-hand side of the table: PP = polyproline, R1-R4 = microtubule binding repeat regions 1-4.

Within the panel, the majority of tau mutants binned into 3 major structure groups: SG1, SG4, and SG9. Collectively, these groups matched at least one replicate for 80% (29/36) of the tau mutant digest profiles. Representative profiles comparing SG1, SG4, and SG9 are shown in Fig. 2E. SG1 which represents the fragment profile for WT tau was the largest grouping of mutants (15/36) and included 10 tau mutants that invariably shared this profile (e.g. A152T, P301S, G335V) and another 5 mutants that generated the profile in at least 1 of 3 independent assays (e.g. G272V, V337M, R406W). The tau mutants belonging to SG1 were spread across the tau sequence, showing no clustering to a specific sequence region. Tau mutants in SG4 (10/36) also did not show any association with a specific location. SG4 contained relatively rare FTLD-tau mutations with the exception of V337M which could generate more than one structure profile (SG1 and SG4). SG9 contained 6 mutants (I260V, G272V, G273R, N279K, L284R, L315R) that did seem to cluster in the first two microtubule binding repeat regions of tau (R1-R2) however, one of the mutants, L315R, was located further downstream in R3. Strikingly, there was little to no overlap between the profiles generated for the tau variants within SG1, SG4, and SG9; V337M was the only mutant shared between major structure groups (SG1 and SG4). With respect to the other structure groups, some contained just a single mutant (G304S, G366R, K369I, P332S, Q336R, E342V, E372G) whereas other groups contained up to 3 mutants. The core packing of aggregates for the vast majority of the mutants we tested have not been previously characterized. However, Aoyagi et al. have compared the packing of WT, P301L and R406W 0N4R tau via pronase treatment^64^. They found that the P301L mutation produced aggregates with a digest fragment profile distinct from WT tau. We also found that P301L tau consistently classified into a different structure group (SG6) than WT (Fig. 2F). The Aoyagi study also reported that WT and R406W tau aggregates shared a similar digest fragment profile. In contrast, we detected that R406W tau aggregate structures could be matched to WT in only two out of three of our trials. For one R406W tau aggregate reaction, we detected a packed core that was distinct from WT. All of the aggregates formed by tau mutants in the panel were confirmed to be positive for ThT binding in our aggregation kinetic assays indicating that they contain beta-sheet rich properties characteristic of amyloids. Thus, our dataset highlights the diversity of amyloid structures formed by these disease-associated mutations and offers evidence of how their pathological misfolding can be readily differentiated from that of WT tau.

One key observation that emerged from our large mutant panel analysis was that a subset of mutants such as V337M generated more than one digest fragment profile between experiments which we termed “variable structure” mutants. Within our panel, 30% (11/36) of the tau mutants assayed had variable structure and are highlighted in Table 1 (red and orange) and in Fig. 2D (orange). These variable structure mutants in general displayed one structure profile that fell into the major structure groups (SG1, SG4, or SG9) and another alternate structure profile. An example of the digest profiles generated by the variable tau mutant G272V are shown in Fig. 2G. The alternate structure could group with other mutant profiles (e.g. P301T, deltaK280 and P301L group in SG6) or could be classified as a unique structure in some cases (e.g. G366R = SG5, K369I = SG7, E342V = SG8). Since we did not detect variable structures for WT and the majority of tau mutants, it suggests that there is a predominant misfolding pathway for most tau variants but that some mutations allow for the occurrence of an early aggregate species or event that can shuttle tau down an alternative divergent misfolding pathway(s).

Even with the large panel of mutants analyzed, no clear relationships could be identified between the biochemical changes in tau caused by mutation and their classification into specific structural groups. No overall trend was observed between the relative locations of the mutations and their grouping into a structure group. The strongest trend was observed for SG9 that contained several mutations clustered between residues 260-284 but mutations within this sequence (deltaK280) did not generate the same structure and mutations outside the 260-284 region could also generate the SG9 fragment profile (L315R). We also found no correlations between changes in the protein’s isoelectric point or hydrophobicity values and structure group classifications (Supp. Fig. 3B-C). We also observed that varying the missense mutation at the same residue (e.g. G335 and P301) could give rise to alternate structure groups. G335V gave a digest profile matching WT tau whereas G335S gave rise to an alternate digest profile (SG2). At the P301 residue, mutation to a leucine (P301L) generated aggregates classed into SG6 (Fig. 2F) whereas mutation to a serine (P301S) gave rise to aggregates similar to WT (SG1). Interestingly, the P301T mutation fell into the variable structure category and could form both SG1 and SG6 structures. These results begin to reveal the complex nature of the effects of tau mutations on aggregate structure formation, where even subtle differences at a given residue can significantly impact the reaction products.

### Aggregation kinetics of individual tau mutants is not a predictor of distinct aggregation structure profiles

We further probed whether the alterations in aggregate structure we detected could be linked to other tau properties such as aggregation kinetics. We carried *in vitro* aggregation kinetic assays for the entire tau mutant panel (WT and 36 mutants). We found that the extracted kinetic parameters (lag time, elongation rate constant, and amplitude) contained some variation when tested across multiple independent experiments and protein preparations as expected (Fig. 3A, Supp. Fig. 5A-B). For WT tau, absolute calculated lag times varied from 1.04 - 2.5 h (median = 1.41 h, quartiles = 1.21, 1.76). The largest variance in lag time was observed for K298E tau (median = 3.84 h, quartiles = 1.89, 6.64). We also noted that large variations in one kinetic parameter for a mutant did not correlate with variability in other kinetic parameters. For example, K298E showed relatively little variability in elongation rate constant or amplitude values compared to most mutants. Within the panel, the P301 mutants (P301L, P301S, P301T) showed the largest variation in elongation rate constant values. There was some agreement between the heparin-induced aggregation kinetic values we obtained and those reported in previous studies for tau mutants. For example, P301L, P301S and P301T mutants had significantly decreased lag times compared to WT^33,46,65,66^. However, we were not able to confirm previous evidence that other tau mutations (G272V, N279K, V337M, R406W) in the context of full-length tau also drove faster aggregation in the presence of heparin^33^ or contrasting slower initial aggregation kinetics reported for R406W^33,66^. In the case of N279K tau, we observed a significantly longer lag time than WT. The reason for this discrepancy is unclear but aggregation kinetics can be modulated by variables such as buffer conditions^67,68^ and the tau isoforms^69^ examined which differed between our current study and previous studies. It is possible that the presence of mutations in the context of other reaction conditions may differentially modulate aggregation kinetics.

**Figure 3.**
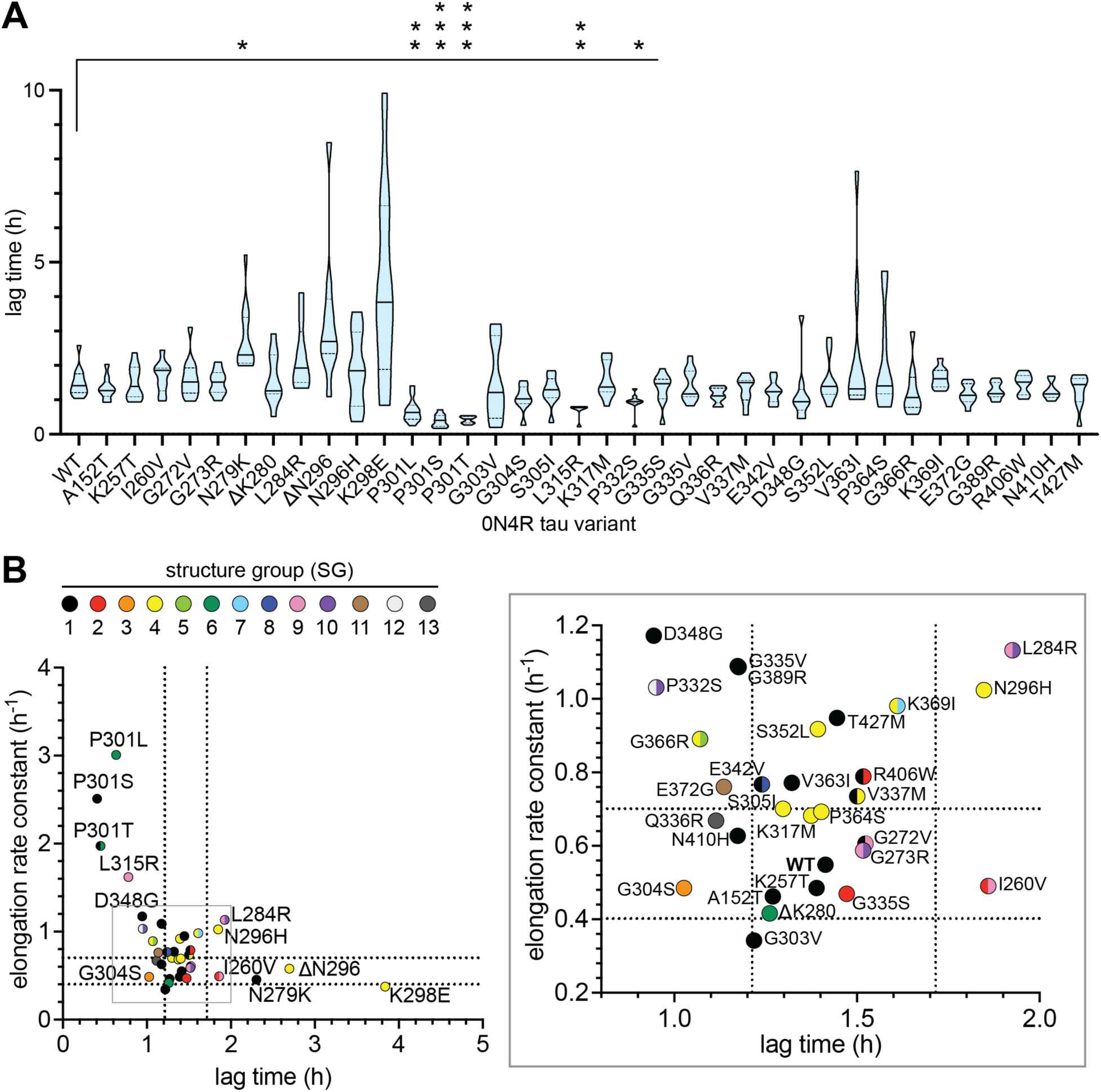
Mutation-dependent alterations in tau aggregation kinetics do not predict changes in aggregate structures. A) Lag time of individual tau WT and mutants. The median (solid line) and quartiles (dotted lines) are plotted for each variant (n ≥ 9 replicates, ≥ 3 independent experiments). A Brown-Forsythe and Welch ANOVA analysis was performed with post-hoc comparisons to the WT tau group. p =* <0.05, **<0.01, ***<0.0001. B) Comparison of WT and tau mutants with respect to: elongation rate constant, lag time, and structure group classification. For each tau variant, median lag time and elongation rate constant values are plotted (n ≥ 9 replicates, ≥ 3 independent experiments). Each tau variant is colored according to its classified structure group as indicated in the legend. Variable structure tau mutants are colored with both corresponding structure groups as half-circles (from n = 3 independent experiments). For WT tau, the quartile boundaries of lag time and elongation rate constant values are marked by dashed lines. Graph showing all assayed tau variants (left) with grey boxed region expanded to aid in comparison of mutants clustered in this region (right).

We next plotted the median values for lag time against elongation rate constant and structure group for all tau variants in the panel (Fig. 3B). We did not observe any clear trends between lag time or elongation rate constant and specific structure groups. For example, tau mutants that promoted the formation of an alternative aggregate structure to WT could still have WT-like aggregation kinetics. Groups of tau mutants that were categorized to have the same aggregate structure profile could have dissimilar aggregation kinetic values in the parameters assessed. We also did not detect a relationship between amplitude and structure groups (Supp. Fig. 5C) or kinetic parameters and variable versus consistent structure mutants (Supp. Fig. 5D). Thus, variable aggregation kinetics can not readily account for the propensity of individual tau mutants to form alternate aggregate structures.

### Examining changes in tau aggregate structure profiles in the context of Cryo-EM tau fibril structures

Current Cryo-EM tau fibril structures contain WT full-length tau isoforms. Thus, we hypothesized that disease-associated mutations identified as forming alternate aggregate structures in our digestion assays may also be more likely to be incompatible with packing in the ordered fibril core. Mutagenesis analysis for disease-associated mutations were carried out in PyMOL on individual solved tau fibril structures for recombinant fibrils and patient-derived fibrils from Alzheimer and CBD samples^10,13,15,70,71^. For each structure, predicted steric clashes induced by mutation are reported in Table 2. Figure 4A maps the location of individual tau mutations and their predicted clashes in the context of the sequences solved in Cryo-EM structures. We first focused on a fibril structure obtained by aggregation of recombinant tau termed the “snake” filament. All mutations predicted to cause a steric clash in our mutational analysis (N279K, deltaK280, L284R, deltaN296, S305I) also produced aggregate structure distinct from WT tau, supporting that these mutations are incompatible with the observed packing in the fibril structure. This was expected given these residues face towards the interface of beta-strand packing interactions (Supp. Fig. 6). However, we also found that the majority of mutations in the known ordered core region (with the exception of P301S and G303V) generated alternate structures to WT tau in our digest assay, despite inducing no detectable steric clash in our computational mutagenesis analysis. Most of the mutated residue sites face outwards from the resolved packed core structure and it is not readily clear how mutation at these sites would induce fibril structure changes (Supp. Fig. 6). Our results could indicate that 1) our WT tau fibril structures are distinct from previously solved structures or 2) that disease associated mutations may act on the misfolding pathway upstream of fibril formation to alter the resulting aggregate structures produced (see discussion). Another key observation is that our mutant panel coverage extends beyond the solved regions for all current tau fibril structures. For example, the recombinant tau fibril structure termed the “snake filament” covers only 44% (16/36) of our examined mutations. Out of the mutations that lie outside the solved structure region, our digest fragment results suggest that 12/20 mutations induce formation of structures distinct from WT tau. This raises the possibility that even though a mutation is not directly within the solved packed core structure, it may still influence core packing through distal interactions or induce local reproducible folding alterations in the more flexible regions of the aggregate core.

**Figure 4.**
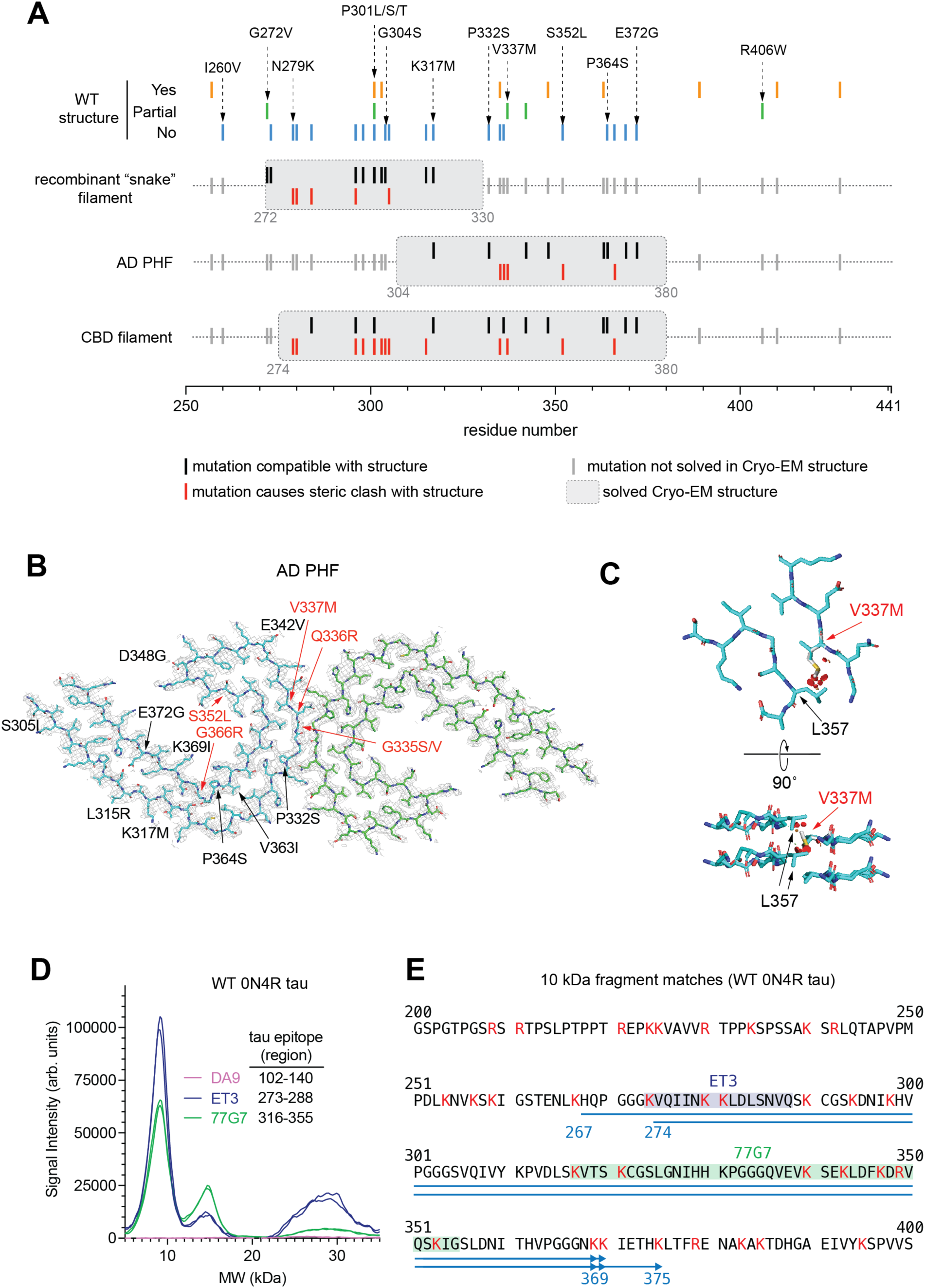
Predicted effects of disease-associated tau mutations in the context of current Cryo-EM tau fibril structures. A) Schematic displaying the relative location of tau mutations with sequence regions of indicated tau fibril structures solved by Cryo-EM. The information in the schematic as well as PDB file information for structures is summarized in Table 2. At the top of the schematic, information is given for each mutation with respect to whether the designated structure group matches with WT (lines: yes=orange, no=blue, partial=green). For each Cryo-EM structure analyzed (grey box highlights solved region), each mutation is colored according to whether it is predicted to cause a steric clash (red line), does not cause a steric clash (black line) or is not solved in the indicated Cryo-EM structure (grey lines). B) Cryo-EM density map (grey mesh) of PHF tau fibril structure from AD patient (PDB: 6HRE). The two individual protofibrils for the solved structure are shown as blue and green backbone traces. Disease associated mutations assayed in our study are labeled on one of the protofibrils (left) according to whether they are predicted to cause a steric clash (red) or not (black) as indicated in A). C) Magnification of residues surrounding V337 (blue backbone) in the same orientation as B) or rotated 90 degrees. The V337M mutation has been superimposed onto the structure (white backbone) and predicted steric clashes with L357 are indicated (red disks). D-E) Epitope mapping of 10 kDa trypsin-resistant fragment for WT 0N4R tau. D) Chromatogram results of capillary electrophoresis immunoassay with the tau antibodies DA9, ET3, and 77G7. The epitope region for each antibody is listed (2N4R numbering). E) Tau amino acid sequence (residues 200-400) with the predicted sequence for the 10 kDa fragment indicated with blue lines (arrow =potential C-terminal ends). The epitope regions for ET3 (blue) and 77G7 (green) antibodies are highlighted on the sequence. Trypsin sites at lysine and arginine residues are indicated as red text.

**Table 2.**
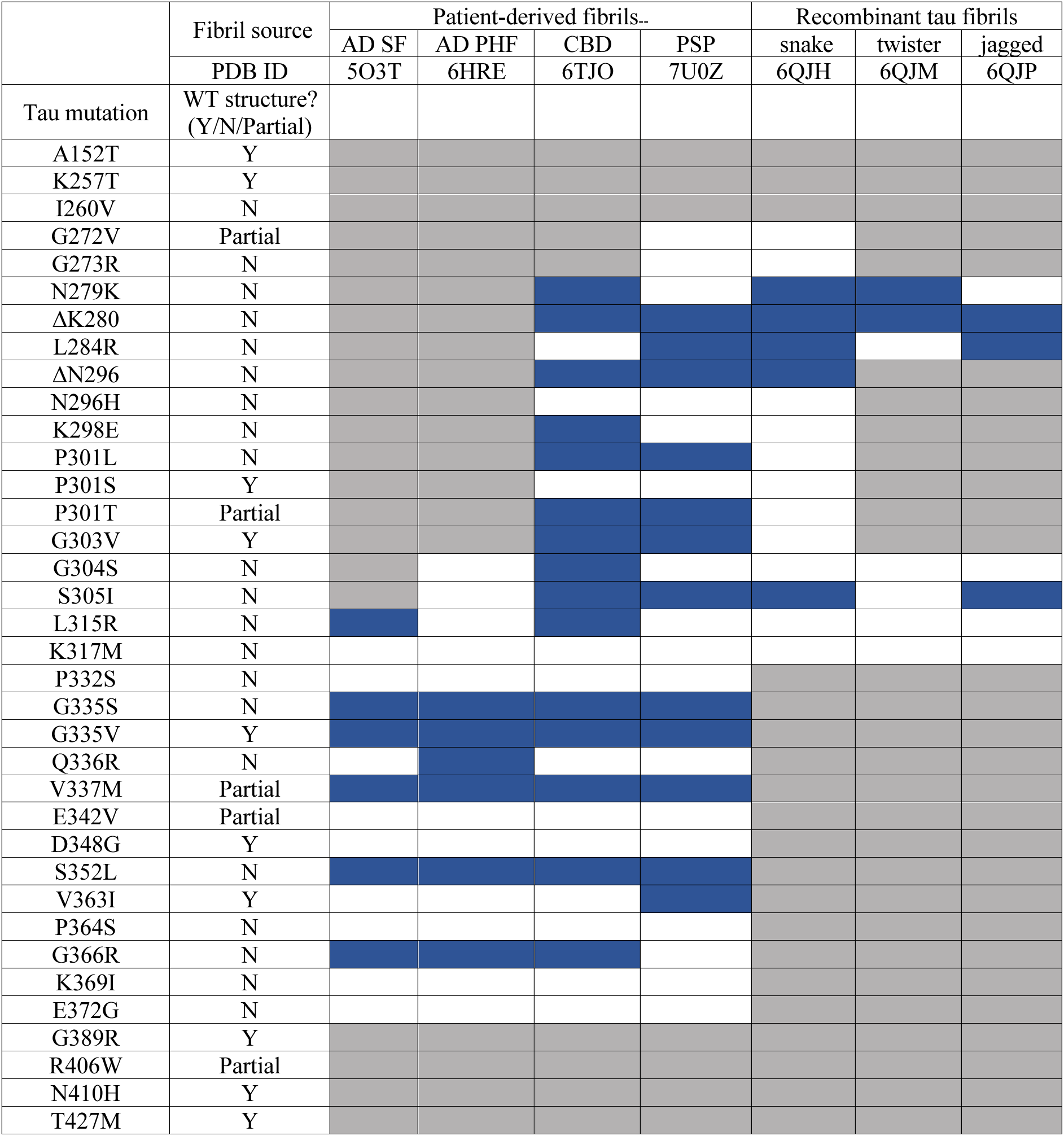
Predicted effect of tau disease-associated mutations on tau fibril structures. Amino acid substitutions were modeled into Cryo-EM resolved tau fibril structures indicated in the table. Using the trypsin digest fragment profiles, mutations binned into the same structure group as WT tau (SG1) versus mutations that bin into another group or only with WT in some trials are labelled as Y(yes), N(no) or Partial, respectively. For each Cryo-EM structure analyzed, cells are colored according to whether the mutations are predicted to cause a steric clash (blue cell) or not (white cells). Mutations that are not resolved in structures are colored as grey cells.

We also carried out an analysis of the compatibility of disease-associated mutations on structures solved for patient-derived fibrils (AD and CBD). In the AD paired helical filament (PHF) structure Q336R, G335S/V, V337M, S352L and G366R were predicted to cause clashes Fig. 4B). Similar to the recombinant tau fibril analysis, we observed all mutations (except G335V) which were predicted to cause clashes did form alternate aggregate structures in our digest assays. As an example, the V337M mutation linked to FTLD-tau in separate 3 families ^72,73^, we predicted it caused potential steric clashes with all patient fibril structures examined: AD, CBD and PSP ^72,73^ (steric clash shown for AD PHF in Fig. 4C). Our digest assays indicate that V337M has the propensity to form non-WT aggregate structures. The finding that V337M and other mutations are not compatible with structures linked to WT tau tauopathies raises the possibility that their pathogenic properties may be due to their inherent propensity to form alternate aggregate structures that promote FTLD-tau specifically.

To gain more insight into the sequences present in the packed core region of our recombinant tau aggregates we performed epitope mapping of trypsin-resistant fragments by capillary gel electrophoresis. We probed WT 0N4R digest fragments with individual tau antibodies (Fig. 4D). Differential positive or negative tau antibody reactivity and apparent MW of individual fragments were used to narrow down the identity of the 10 kDa fragment to five potential sequence regions (Fig. 4E). The five predicted sequences largely overlap with the most N-terminal sequence fragment covering residues 267-369 and the most C-terminal fragment covering residues 274-375. The finding that this 10 kDa fragment is consistent in our trypsin digest profile as one of the 3 major trypsin-resistant bands produced suggests that it is a main component of the protected core region of our aggregates. The predicted sequence coverage indicates that it is larger than the region solved by Cryo-EM for recombinant tau aggregates (272-330). The 10 kDa fragment more closely aligns with trypsin-resistant core sequences identified for patient-derived fibrils in CBD (268-369/387/395), PSP (260/268-395) or genetic FTLD-tau (268-369/387/395)^7^. It also overlaps with highly GluC-resistant sequence regions (223-380) reported for 1N4R recombinant tau fibrils^6^.

## Discussion

There remains much to be learned about how tau is able to misfold and aggregate into the multiple amyloid core confirmations observed in tauopathies and how tau’s primary sequence contributes to specific misfolding events. Our current study takes advantage of missense mutations in tau linked to tauopathies to shed insight into the formation of specific tau aggregate structures. We reveal that single missense mutations introduced into human full-length 0N4R tau are sufficient to modulate its misfolding pathway *in vitro*, leading to the production of alternate aggregate structures. Using the high-throughput tau purification platform we developed allowed us to perform direct comparisons of 36 tau mutants and WT tau with respect to their aggregate properties: structure and kinetics. We demonstrate the ability of a subset of tau mutations to inherently modulate tau aggregate structure formation, defining 13 distinct tau aggregate structure profiles within our panel. Our results offer a view into the complexity of the relationship between tau primary sequence and aggregation and provides a roadmap with which to systematically begin to dissect the fundamental principles underlying tau misfolding and aggregation.

Several atomic resolution structures have been solved for tau fibrils derived from tauopathies. However, the ability to perform similar studies in the case of familial tau mutations linked to tauopathies are limited by the extremely rare incidence of these mutations in general, with some mutations being reported in as few as 2 cases^74,75^ (e.g. deltaK280). Our *in vitro* methods with recombinant tau protein provides an opportunity to directly test how tau mutations modulate aggregate structure formation under controlled and consistent conditions. The current study specifically examined the effects of tau mutations in the context of the 0N4R tau isoform. The results form a foundation in which to test other variables that may influence tau aggregate structure formation. For example, six tau isoforms are normally expressed in the neurons of the adult CNS^76^ and it will be interesting to test if tau mutation effects on aggregate structure are applicable across all isoforms or behave similarly in aggregation reactions where multiple tau isoforms are present. Individual disease-associated tau mutations have been shown to cause tau aggregate pathology that consists primarily of 3R, 4R, or a mixture of tau isoforms^29,41^. In some cases, the inclusion of some tau isoforms but not others in aggregates may be caused by preferential expression of isoforms via mechanisms such as differential splicing^30–32^. However, another possibility is that some tau mutations drive isoform-specific promotion of pathogenic tau aggregate structures or ineffectively template some tau isoforms. This is plausible given that another group has already found differences in the packed core conformations between 1N3R and 1N4R heparin-induced aggregates including an additional protease protected C-terminal sequence region for 1N3R^6^. Tau aggregate structures can also be modulated by the accelerants^77^ or buffer conditions^78^ of the aggregation reaction. The packed fibril core of tau fibrils show distinct conformations when aggregation is initiated by heparin^70^, RNA^78,79^ or phosphoserine^78^. Using our assays, we can begin to tease out the relative contributions of tau primary sequence or other modifications (e.g. PTMs) versus added cofactors which may offer additional insight into how tau aggregation can be triggered in living cells.

We chose trypsin to identify changes in the protease-resistance of tau aggregates because the high incidence of regularly spaced lysine and arginine residues throughout the tau sequence (Supp. Fig. 3A) made it a sensitive indicator of structure changes that could be easily mapped. A potential caveat of our trypsin digest experiments is that multiple disease associated mutations involve changes to or from lysine(K) and arginine(R) residues. Thus, it is possible that new aggregate structure profiles could be the result of gain or loss of a trypsin cleavage site. However, our data does not support this as being a general effect of K/R mutations. First, multiple mutations spread across different locations in tau (e.g. K298E, K317M, G366R, K369I) can produce the same trypsin-resistant profile boned as structure group 4 (SG4). Second, mutations such as L315R and Q336R introduce a new tryptic site located in the middle of the sequence of all available Cryo-EM structures as well as the 10 kDa fragment (residues within 267-375) identified in our WT tau digestion profile. Yet, a banding pattern reflective of trypsin digestion at this newly introduced site which would essentially cleave the 10 kDa band into two smaller fragments, is not observed (Supp. Fig. 7). Thus, we contend that tau mutants in our study involving K or R mutations can modulate tau aggregate core packing leading to the observed changes in protease-resistant profiles.

An unexpected finding from our study is that the effects of mutations on aggregate structures are not readily reconciled with current atomic resolution structure data on recombinant tau fibrils^70^. Several tau mutations that were predicted to be compatible with the fibril core solved by Cryo-EM generated structures distinguishable from WT. Moreover, tau mutations outside the region of the solved core structure were also found to alter aggregate structure formation. There could be several reasons to explain these discrepancies. Cryo-EM structures of recombinant tau induced in the presence of heparin use 2N4R tau^70^. It is possible that the specific tau isoform or other reaction conditions used in the previous study leads to an aggregate structure distinct from those generated by our conditions. Another possibility is that tau mutations alter upstream misfolding events that shuttle the misfolding pathway towards the production of alternate aggregate structures. In the case of tau mutations located outside the solved packed core region, we also postulate that mutation effects could be due to changes in the packing in more open/flexible structured regions in fibrils. For WT 0N4R tau, we mapped our 10 kDa fragment to a region within residues 267-375. All but two of the tau mutations that altered aggregate structure (I260V, R406W) in this study are contained within this region. Thus, our results offer new insights into how tau’s primary sequence may contribute to shaping tau aggregate structure in regions not yet resolvable by techniques such as Cryo-EM.

Using our new high-throughput methods for tau purification and our trypsin digest assay for comparing aggregate structures, we hope to assist in defining the complex relationships between tau sequence, aggregate properties and the biological effects of aggregates at the cellular and *in vivo* levels. Multiple factors can contribute to the pathogenicity of a disease-associated tau mutation. Tau mutations such as P301L enhance the propensity of tau to aggregate however, this is not the case for other tau mutations linked to tauopathy. Our results open up the possibility that tau mutations with WT-like or slow aggregation kinetics may be pathogenic, in part, via their ability to assemble into alternate tau aggregate structures. Future characterization comparing the ability of distinct tau aggregate structures to promote seeding, propagation, and evade protein quality control factors will dissect the fundamental features of tau aggregates responsible for their induced pathobiology.

## Acknowledgements

We thank G. Eskandari-Sedighi and A. Castle for advice regarding capillary gel electrophoresis, and J. Moore of the Alberta Proteomics and Mass Spectrometry Facility (University of Alberta) for assistance with mass spectrometry sample preparation and data analysis. We thank Peter Davies for generously providing tau antibodies used in this study. We thank R. Fahlman and V. Sim for critical discussions of the project during its development. This work was funded by grants to S.A.M from Alberta Innovates, the Alzheimer’s Society of Alberta and Northwest Territories, NSERC, and donors of the ADR; a program of the BrightFocus Foundation (Award #: A2022044S).

## Author Contributions

K.S., T.P., J.H., A.Y. and S.A.M. conceived and designed the study. K.S., T.P., S.K., J.H., O.J. and A.Y. acquired, analyzed, or interpreted data. K.S., T.P., S.K., J.H. and S.A.M. drafted and revised the manuscript.

## References

(1) Kovacs, G. G. Invited Review: Neuropathology of Tauopathies: Principles and Practice. Neuropathol. Appl. Neurobiol. 2015, 41 (1), 3–23. https://doi.org/10.1111/nan.12208.

(2) Lee, V. M.-Y.; Goedert, M.; Trojanowski, J. Q. Neurodegenerative Tauopathies. Annu. Rev. Neurosci. 2001, 24 (1), 1121–1159. https://doi.org/10.1146/annurev.neuro.24.1.1121.

(3) Wegmann, S.; Medalsy, I. D.; Mandelkow, E.; Müller, D. J. The Fuzzy Coat of Pathological Human Tau Fibrils Is a Two-Layered Polyelectrolyte Brush. Proc. Natl. Acad. Sci. U. S. A. 2013, 110 (4), E313–321. https://doi.org/10.1073/pnas.1212100110.

(4) Barghorn, S.; Davies, P.; Mandelkow, E. Tau Paired Helical Filaments from Alzheimer’s Disease Brain and Assembled in Vitro Are Based on Beta-Structure in the Core Domain. Biochemistry 2004, 43 (6), 1694–1703. https://doi.org/10.1021/bi0357006.

(5) Wischik, C. M.; Novak, M.; Edwards, P. C.; Klug, A.; Tichelaar, W.; Crowther, R. A. Structural Characterization of the Core of the Paired Helical Filament of Alzheimer Disease. Proc. Natl. Acad. Sci. U. S. A. 1988, 85 (13), 4884–4888. https://doi.org/10.1073/pnas.85.13.4884.

(6) Caroux, E.; Redeker, V.; Madiona, K.; Melki, R. Structural Mapping Techniques Distinguish the Surfaces of Fibrillar 1N3R and 1N4R Human Tau. J. Biol. Chem. 2021, 297 (5), 101252. https://doi.org/10.1016/j.jbc.2021.101252.

(7) Taniguchi-Watanabe, S.; Arai, T.; Kametani, F.; Nonaka, T.; Masuda-Suzukake, M.; Tarutani, A.; Murayama, S.; Saito, Y.; Arima, K.; Yoshida, M.; Akiyama, H.; Robinson, A.; Mann, D. M. A.; Iwatsubo, T.; Hasegawa, M. Biochemical Classification of Tauopathies by Immunoblot, Protein Sequence and Mass Spectrometric Analyses of Sarkosyl-Insoluble and Trypsin-Resistant Tau. Acta Neuropathol. (Berl.) 2016, 131 (2), 267–280. https://doi.org/10.1007/s00401-015-1503-3.

(8) Hasegawa, M.; Watanabe, S.; Kondo, H.; Akiyama, H.; Mann, D. M. A.; Saito, Y.; Murayama, S. 3R and 4R Tau Isoforms in Paired Helical Filaments in Alzheimer’s Disease. Acta Neuropathol. (Berl.) 2014, 127 (2), 303–305. https://doi.org/10.1007/s00401-013-1191-9.

(9) Jakes, R.; Novak, M.; Davison, M.; Wischik, C. M. Identification of 3- and 4-Repeat Tau Isoforms within the PHF in Alzheimer’s Disease. EMBO J. 1991, 10 (10), 2725–2729. https://doi.org/10.1002/j.1460-2075.1991.tb07820.x.

(10) Falcon, B.; Zhang, W.; Schweighauser, M.; Murzin, A. G.; Vidal, R.; Garringer, H. J.; Ghetti, B.; Scheres, S. H. W.; Goedert, M. Tau Filaments from Multiple Cases of Sporadic and Inherited Alzheimer’s Disease Adopt a Common Fold. Acta Neuropathol. (Berl.) 2018, 136 (5), 699–708. https://doi.org/10.1007/s00401-018-1914-z.

(11) Falcon, B.; Zhang, W.; Murzin, A. G.; Murshudov, G.; Garringer, H. J.; Vidal, R.; Crowther, R. A.; Ghetti, B.; Scheres, S. H. W.; Goedert, M. Structures of Filaments from Pick’s Disease Reveal a Novel Tau Protein Fold. Nature 2018, 561 (7721), 137–140. https://doi.org/10.1038/s41586-018-0454-y.

(12) Falcon, B.; Zivanov, J.; Zhang, W.; Murzin, A. G.; Garringer, H. J.; Vidal, R.; Crowther, R. A.; Newell, K. L.; Ghetti, B.; Goedert, M.; Scheres, S. H. W. Novel Tau Filament Fold in Chronic Traumatic Encephalopathy Encloses Hydrophobic Molecules. Nature 2019, 568 (7752), 420–423. https://doi.org/10.1038/s41586-019-1026-5.

(13) Zhang, W.; Tarutani, A.; Newell, K. L.; Murzin, A. G.; Matsubara, T.; Falcon, B.; Vidal, R.; Garringer, H. J.; Shi, Y.; Ikeuchi, T.; Murayama, S.; Ghetti, B.; Hasegawa, M.; Goedert, M.; Scheres, S. H. W. Novel Tau Filament Fold in Corticobasal Degeneration. Nature 2020, 580 (7802), 283–287. https://doi.org/10.1038/s41586-020-2043-0.

(14) Shi, Y.; Zhang, W.; Yang, Y.; Murzin, A. G.; Falcon, B.; Kotecha, A.; van Beers, M.; Tarutani, A.; Kametani, F.; Garringer, H. J.; Vidal, R.; Hallinan, G. I.; Lashley, T.; Saito, Y.; Murayama, S.; Yoshida, M.; Tanaka, H.; Kakita, A.; Ikeuchi, T.; Robinson, A. C.; Mann, D. M. A.; Kovacs, G. G.; Revesz, T.; Ghetti, B.; Hasegawa, M.; Goedert, M.; Scheres, S. H. W. Structure-Based Classification of Tauopathies. Nature 2021, 598 (7880), 359–363. https://doi.org/10.1038/s41586-021-03911-7.

(15) Fitzpatrick, A. W. P.; Falcon, B.; He, S.; Murzin, A. G.; Murshudov, G.; Garringer, H. J.; Crowther, R. A.; Ghetti, B.; Goedert, M.; Scheres, S. H. W. Cryo-EM Structures of Tau Filaments from Alzheimer’s Disease. Nature 2017, 547 (7662), 185–190. https://doi.org/10.1038/nature23002.

(16) Frost, B.; Diamond, M. I. Prion-like Mechanisms in Neurodegenerative Diseases. Nat. Rev. Neurosci. 2010, 11 (3), 155–159. https://doi.org/10.1038/nrn2786.

(17) Dujardin, S.; Hyman, B. T. Tau Prion-Like Propagation: State of the Art and Current Challenges. In *Tau Biology*; Takashima, A., Wolozin, B., Buee, L., Eds.; Advances in Experimental Medicine and Biology; Springer Singapore: Singapore, 2019; Vol. 1184, pp 305–325. https://doi.org/10.1007/978-981-32-9358-8_23.

(18) Clavaguera, F.; Bolmont, T.; Crowther, R. A.; Abramowski, D.; Frank, S.; Probst, A.; Fraser, G.; Stalder, A. K.; Beibel, M.; Staufenbiel, M.; Jucker, M.; Goedert, M.; Tolnay, M. Transmission and Spreading of Tauopathy in Transgenic Mouse Brain. Nat. Cell Biol. 2009, 11 (7), 909–913. https://doi.org/10.1038/ncb1901.

(19) Sanders, D. W.; Kaufman, S. K.; DeVos, S. L.; Sharma, A. M.; Mirbaha, H.; Li, A.; Barker, S. J.; Foley, A. C.; Thorpe, J. R.; Serpell, L. C.; Miller, T. M.; Grinberg, L. T.; Seeley, W. W.; Diamond, M. I. Distinct Tau Prion Strains Propagate in Cells and Mice and Define Different Tauopathies. Neuron 2014, 82 (6), 1271–1288. https://doi.org/10.1016/j.neuron.2014.04.047.

(20) Kaufman, S. K.; Sanders, D. W.; Thomas, T. L.; Ruchinskas, A. J.; Vaquer-Alicea, J.; Sharma, A. M.; Miller, T. M.; Diamond, M. I. Tau Prion Strains Dictate Patterns of Cell Pathology, Progression Rate, and Regional Vulnerability In Vivo. Neuron 2016, 92 (4), 796–812. https://doi.org/10.1016/j.neuron.2016.09.055.

(21) Guo, J. L.; Narasimhan, S.; Changolkar, L.; He, Z.; Stieber, A.; Zhang, B.; Gathagan, R. J.; Iba, M.; McBride, J. D.; Trojanowski, J. Q.; Lee, V. M. Y. Unique Pathological Tau Conformers from Alzheimer’s Brains Transmit Tau Pathology in Nontransgenic Mice. J. Exp. Med. 2016, 213 (12), 2635–2654. https://doi.org/10.1084/jem.20160833.

(22) Woerman, A. L.; Aoyagi, A.; Patel, S.; Kazmi, S. A.; Lobach, I.; Grinberg, L. T.; McKee, A. C.; Seeley, W. W.; Olson, S. H.; Prusiner, S. B. Tau Prions from Alzheimer’s Disease and Chronic Traumatic Encephalopathy Patients Propagate in Cultured Cells. Proc. Natl. Acad. Sci. U. S. A. 2016, 113 (50), E8187–E8196. https://doi.org/10.1073/pnas.1616344113.

(23) Narasimhan, S.; Guo, J. L.; Changolkar, L.; Stieber, A.; McBride, J. D.; Silva, L. V.; He, Z.; Zhang, B.; Gathagan, R. J.; Trojanowski, J. Q.; Lee, V. M. Y. Pathological Tau Strains from Human Brains Recapitulate the Diversity of Tauopathies in Nontransgenic Mouse Brain. J. Neurosci. Off. J. Soc. Neurosci. 2017, 37 (47), 11406–11423. https://doi.org/10.1523/JNEUROSCI.1230-17.2017.

(24) Ghetti, B.; Oblak, A. L.; Boeve, B. F.; Johnson, K. A.; Dickerson, B. C.; Goedert, M. Invited Review: Frontotemporal Dementia Caused by Microtubule-Associated Protein Tau Gene (MAPT) Mutations: A Chameleon for Neuropathology and Neuroimaging. Neuropathol. Appl. Neurobiol. 2015, 41 (1), 24–46. https://doi.org/10.1111/nan.12213.

(25) Pottier, C.; Ravenscroft, T. A.; Sanchez-Contreras, M.; Rademakers, R. Genetics of FTLD: Overview and What Else We Can Expect from Genetic Studies. J. Neurochem. 2016, 138 *Suppl 1*, 32–53. https://doi.org/10.1111/jnc.13622.

(26) Forrest, S. L.; Halliday, G. M.; McCann, H.; McGeachie, A. B.; McGinley, C. V.; Hodges, J. R.; Piguet, O.; Kwok, J. B.; Spillantini, M. G.; Kril, J. J. Heritability in Frontotemporal Tauopathies. Alzheimers Dement. Amst. Neth. 2019, 11, 115–124. https://doi.org/10.1016/j.dadm.2018.12.001.

(27) Moore, K. M.; Nicholas, J.; Grossman, M.; McMillan, C. T.; Irwin, D. J.; Massimo, L.; Van Deerlin, V. M.; Warren, J. D.; Fox, N. C.; Rossor, M. N.; Mead, S.; Bocchetta, M.; Boeve, B. F.; Knopman, D. S.; Graff-Radford, N. R.; Forsberg, L. K.; Rademakers, R.; Wszolek, Z. K.; van Swieten, J. C.; Jiskoot, L. C.; Meeter, L. H.; Dopper, E. G.; Papma, J. M.; Snowden, J. S.; Saxon, J.; Jones, M.; Pickering-Brown, S.; Le Ber, I.; Camuzat, A.; Brice, A.; Caroppo, P.; Ghidoni, R.; Pievani, M.; Benussi, L.; Binetti, G.; Dickerson, B. C.; Lucente, D.; Krivensky, S.; Graff, C.; Öijerstedt, L.; Fallström, M.; Thonberg, H.; Ghoshal, N.; Morris, J. C.; Borroni, B.; Benussi, A.; Padovani, A.; Galimberti, D.; Scarpini, E.; Fumagalli, G. G.; Mackenzie, I. R.; Hsiung, G.-Y. R.; Sengdy, P.; Boxer, A. L.; Rosen, H.; Taylor, J. B.; Synofzik, M.; Wilke, C.; Sulzer, P.; Hodges, J. R.; Halliday, G.; Kwok, J.; Sanchez-Valle, R.; Lladó, A.; Borrego-Ecija, S.; Santana, I.; Almeida, M. R.; Tábuas-Pereira, M.; Moreno, F.; Barandiaran, M.; Indakoetxea, B.; Levin, J.; Danek, A.; Rowe, J. B.; Cope, T. E.; Otto, M.; Anderl-Straub, S.; de Mendonça, A.; Maruta, C.; Masellis, M.; Black, S. E.; Couratier, P.; Lautrette, G.; Huey, E. D.; Sorbi, S.; Nacmias, B.; Laforce, R.; Tremblay, M.-P. L.; Vandenberghe, R.; Damme, P. V.; Rogalski, E. J.; Weintraub, S.; Gerhard, A.; Onyike, C. U.; Ducharme, S.; Papageorgiou, S. G.; Ng, A. S. L.; Brodtmann, A.; Finger, E.; Guerreiro, R.; Bras, J.; Rohrer, J. D.; FTD Prevention Initiative. Age at Symptom Onset and Death and Disease Duration in Genetic Frontotemporal Dementia: An International Retrospective Cohort Study. Lancet Neurol. 2020, 19 (2), 145–156. https://doi.org/10.1016/S1474-4422(19)30394-1.

(28) Young, A. L.; Bocchetta, M.; Russell, L. L.; Convery, R. S.; Peakman, G.; Todd, E.; Cash, D. M.; Greaves, C. V.; van Swieten, J.; Jiskoot, L.; Seelaar, H.; Moreno, F.; Sanchez-Valle, R.; Borroni, B.; Laforce, R.; Masellis, M.; Tartaglia, M. C.; Graff, C.; Galimberti, D.; Rowe, J. B.; Finger, E.; Synofzik, M.; Vandenberghe, R.; de Mendonça, A.; Tagliavini, F.; Santana, I.; Ducharme, S.; Butler, C.; Gerhard, A.; Levin, J.; Danek, A.; Otto, M.; Sorbi, S.; Williams, S. C. R.; Alexander, D. C.; Rohrer, J. D.; Genetic FTD Initiative (GENFI). Characterizing the Clinical Features and Atrophy Patterns of MAPT-Related Frontotemporal Dementia With Disease Progression Modeling. Neurology 2021, 97 (9), e941–e952. https://doi.org/10.1212/WNL.0000000000012410.

(29) Forrest, S. L.; Kril, J. J.; Stevens, C. H.; Kwok, J. B.; Hallupp, M.; Kim, W. S.; Huang, Y.; McGinley, C. V.; Werka, H.; Kiernan, M. C.; Götz, J.; Spillantini, M. G.; Hodges, J. R.; Ittner, L. M.; Halliday, G. M. Retiring the Term FTDP-17 as MAPT Mutations Are Genetic Forms of Sporadic Frontotemporal Tauopathies. Brain J. Neurol. 2018, 141 (2), 521–534. https://doi.org/10.1093/brain/awx328.

(30) Varani, L.; Hasegawa, M.; Spillantini, M. G.; Smith, M. J.; Murrell, J. R.; Ghetti, B.; Klug, A.; Goedert, M.; Varani, G. Structure of Tau Exon 10 Splicing Regulatory Element RNA and Destabilization by Mutations of Frontotemporal Dementia and Parkinsonism Linked to Chromosome 17. Proc. Natl. Acad. Sci. U. S. A. 1999, 96 (14), 8229–8234. https://doi.org/10.1073/pnas.96.14.8229.

(31) D’Souza, I.; Poorkaj, P.; Hong, M.; Nochlin, D.; Lee, V. M.; Bird, T. D.; Schellenberg, G. D. Missense and Silent Tau Gene Mutations Cause Frontotemporal Dementia with Parkinsonism-Chromosome 17 Type, by Affecting Multiple Alternative RNA Splicing Regulatory Elements. Proc. Natl. Acad. Sci. U. S. A. 1999, 96 (10), 5598–5603. https://doi.org/10.1073/pnas.96.10.5598.

(32) Hasegawa, M.; Smith, M. J.; Iijima, M.; Tabira, T.; Goedert, M. FTDP-17 Mutations N279K and S305N in Tau Produce Increased Splicing of Exon 10. FEBS Lett. 1999, 443 (2), 93–96. https://doi.org/10.1016/s0014-5793(98)01696-2.

(33) Barghorn, S.; Zheng-Fischhöfer, Q.; Ackmann, M.; Biernat, J.; von Bergen, M.; Mandelkow, E. M.; Mandelkow, E. Structure, Microtubule Interactions, and Paired Helical Filament Aggregation by Tau Mutants of Frontotemporal Dementias. Biochemistry 2000, 39 (38), 11714–11721. https://doi.org/10.1021/bi000850r.

(34) Goedert, M.; Jakes, R.; Crowther, R. A. Effects of Frontotemporal Dementia FTDP-17 Mutations on Heparin-Induced Assembly of Tau Filaments. FEBS Lett. 1999, 450 (3), 306–311. https://doi.org/10.1016/s0014-5793(99)00508-6.

(35) Combs, B.; Gamblin, T. C. FTDP-17 Tau Mutations Induce Distinct Effects on Aggregation and Microtubule Interactions. Biochemistry 2012, 51 (43), 8597–8607. https://doi.org/10.1021/bi3010818.

(36) Kfoury, N.; Holmes, B. B.; Jiang, H.; Holtzman, D. M.; Diamond, M. I. Trans-Cellular Propagation of Tau Aggregation by Fibrillar Species. J. Biol. Chem. 2012, 287 (23), 19440– 19451. https://doi.org/10.1074/jbc.M112.346072.

(37) Strang, K. H.; Croft, C. L.; Sorrentino, Z. A.; Chakrabarty, P.; Golde, T. E.; Giasson, B. I. Distinct Differences in Prion-like Seeding and Aggregation between Tau Protein Variants Provide Mechanistic Insights into Tauopathies. J. Biol. Chem. 2018, 293 (7), 2408–2421. https://doi.org/10.1074/jbc.M117.815357.

(38) Fischer, D.; Mukrasch, M. D.; von Bergen, M.; Klos-Witkowska, A.; Biernat, J.; Griesinger, C.; Mandelkow, E.; Zweckstetter, M. Structural and Microtubule Binding Properties of Tau Mutants of Frontotemporal Dementias. Biochemistry 2007, 46 (10), 2574–2582. https://doi.org/10.1021/bi061318s.

(39) LeBoeuf, A. C.; Levy, S. F.; Gaylord, M.; Bhattacharya, A.; Singh, A. K.; Jordan, M. A.; Wilson, L.; Feinstein, S. C. FTDP-17 Mutations in Tau Alter the Regulation of Microtubule Dynamics: An “Alternative Core” Model for Normal and Pathological Tau Action. J. Biol. Chem. 2008, 283 (52), 36406–36415. https://doi.org/10.1074/jbc.M803519200.

(40) Xia, Y.; Sorrentino, Z. A.; Kim, J. D.; Strang, K. H.; Riffe, C. J.; Giasson, B. I. Impaired Tau-Microtubule Interactions Are Prevalent among Pathogenic Tau Variants Arising from Missense Mutations. J. Biol. Chem. 2019, 294 (48), 18488–18503. https://doi.org/10.1074/jbc.RA119.010178.

(41) Hong, M.; Zhukareva, V.; Vogelsberg-Ragaglia, V.; Wszolek, Z.; Reed, L.; Miller, B. I.; Geschwind, D. H.; Bird, T. D.; McKeel, D.; Goate, A.; Morris, J. C.; Wilhelmsen, K. C.; Schellenberg, G. D.; Trojanowski, J. Q.; Lee, V. M. Mutation-Specific Functional Impairments in Distinct Tau Isoforms of Hereditary FTDP-17. Science 1998, 282 (5395), 1914–1917. https://doi.org/10.1126/science.282.5395.1914.

(42) Gunawardana, C. G.; Mehrabian, M.; Wang, X.; Mueller, I.; Lubambo, I. B.; Jonkman, J. E. N.; Wang, H.; Schmitt-Ulms, G. The Human Tau Interactome: Binding to the Ribonucleoproteome, and Impaired Binding of the Proline-to-Leucine Mutant at Position 301 (P301L) to Chaperones and the Proteasome. Mol. Cell. Proteomics MCP 2015, 14 (11), 3000–3014. https://doi.org/10.1074/mcp.M115.050724.

(43) Tracy, T. E.; Madero-Pérez, J.; Swaney, D. L.; Chang, T. S.; Moritz, M.; Konrad, C.; Ward, M. E.; Stevenson, E.; Hüttenhain, R.; Kauwe, G.; Mercedes, M.; Sweetland-Martin, L.; Chen, X.; Mok, S.-A.; Wong, M. Y.; Telpoukhovskaia, M.; Min, S.-W.; Wang, C.; Sohn, P. D.; Martin, J.; Zhou, Y.; Luo, W.; Trojanowski, J. Q.; Lee, V. M. Y.; Gong, S.; Manfredi, G.; Coppola, G.; Krogan, N. J.; Geschwind, D. H.; Gan, L. Tau Interactome Maps Synaptic and Mitochondrial Processes Associated with Neurodegeneration. Cell 2022, 185 (4), 712–728.e14. https://doi.org/10.1016/j.cell.2021.12.041.

(44) Dayanandan, R.; Van Slegtenhorst, M.; Mack, T. G.; Ko, L.; Yen, S. H.; Leroy, K.; Brion, J. P.; Anderton, B. H.; Hutton, M.; Lovestone, S. Mutations in Tau Reduce Its Microtubule Binding Properties in Intact Cells and Affect Its Phosphorylation. FEBS Lett. 1999, 446 (2–3), 228–232. https://doi.org/10.1016/s0014-5793(99)00222-7.

(45) Kimura, T.; Hosokawa, T.; Taoka, M.; Tsutsumi, K.; Ando, K.; Ishiguro, K.; Hosokawa, M.; Hasegawa, M.; Hisanaga, S.-I. Quantitative and Combinatory Determination of in Situ Phosphorylation of Tau and Its FTDP-17 Mutants. Sci. Rep. 2016, 6, 33479. https://doi.org/10.1038/srep33479.

(46) Mok, S.-A.; Condello, C.; Freilich, R.; Gillies, A.; Arhar, T.; Oroz, J.; Kadavath, H.; Julien, O.; Assimon, V. A.; Rauch, J. N.; Dunyak, B. M.; Lee, J.; Tsai, F. T. F.; Wilson, M. R.; Zweckstetter, M.; Dickey, C. A.; Gestwicki, J. E. Mapping Interactions with the Chaperone Network Reveals Factors That Protect against Tau Aggregation. Nat. Struct. Mol. Biol. 2018, 25 (5), 384–393. https://doi.org/10.1038/s41594-018-0057-1.

(47) Caballero, B.; Wang, Y.; Diaz, A.; Tasset, I.; Juste, Y. R.; Stiller, B.; Mandelkow, E.-M.; Mandelkow, E.; Cuervo, A. M. Interplay of Pathogenic Forms of Human Tau with Different Autophagic Pathways. Aging Cell 2018, 17 (1), e12692. https://doi.org/10.1111/acel.12692.

(48) Mahali, S.; Martinez, R.; King, M.; Verbeck, A.; Harari, O.; Benitez, B. A.; Horie, K.; Sato, C.; Temple, S.; Karch, C. M. Defective Proteostasis in Induced Pluripotent Stem Cell Models of Frontotemporal Lobar Degeneration. Transl. Psychiatry 2022, 12 (1), 508. https://doi.org/10.1038/s41398-022-02274-5.

(49) Sohn, P. D.; Huang, C. T.-L.; Yan, R.; Fan, L.; Tracy, T. E.; Camargo, C. M.; Montgomery, K. M.; Arhar, T.; Mok, S.-A.; Freilich, R.; Baik, J.; He, M.; Gong, S.; Roberson, E. D.; Karch, C. M.; Gestwicki, J. E.; Xu, K.; Kosik, K. S.; Gan, L. Pathogenic Tau Impairs Axon Initial Segment Plasticity and Excitability Homeostasis. Neuron 2019, 104 (3), 458–470.e5. https://doi.org/10.1016/j.neuron.2019.08.008.

(50) Combs, B.; Christensen, K. R.; Richards, C.; Kneynsberg, A.; Mueller, R. L.; Morris, S. L.; Morfini, G. A.; Brady, S. T.; Kanaan, N. M. Frontotemporal Lobar Dementia Mutant Tau Impairs Axonal Transport through a Protein Phosphatase 1γ-Dependent Mechanism. J. Neurosci. Off. J. Soc. Neurosci. 2021, 41 (45), 9431–9451. https://doi.org/10.1523/JNEUROSCI.1914-20.2021.

(51) Denk, F.; Wade-Martins, R. Knock-out and Transgenic Mouse Models of Tauopathies. Neurobiol. Aging 2009, 30 (1), 1–13. https://doi.org/10.1016/j.neurobiolaging.2007.05.010.

(52) Frost, B.; Ollesch, J.; Wille, H.; Diamond, M. I. Conformational Diversity of Wild-Type Tau Fibrils Specified by Templated Conformation Change. J. Biol. Chem. 2009, 284 (6), 3546–3551. https://doi.org/10.1074/jbc.M805627200.

(53) Chen, D.; Bali, S.; Singh, R.; Wosztyl, A.; Mullapudi, V.; Vaquer-Alicea, J.; Jayan, P.; Melhem, S.; Seelaar, H.; van Swieten, J. C.; Diamond, M. I.; Joachimiak, L. A. FTD-Tau S320F Mutation Stabilizes Local Structure and Allosterically Promotes Amyloid Motif-Dependent Aggregation. Nat. Commun. 2023, 14 (1), 1625. https://doi.org/10.1038/s41467-023-37274-6.

(54) Moore, C. L.; Huang, M. H.; Robbennolt, S. A.; Voss, K. R.; Combs, B.; Gamblin, T. C.; Goux, W. J. Secondary Nucleating Sequences Affect Kinetics and Thermodynamics of Tau Aggregation. Biochemistry 2011, 50 (50), 10876–10886. https://doi.org/10.1021/bi2014745.

(55) Winsor, C. P. The Gompertz Curve as a Growth Curve. Proc. Natl. Acad. Sci. U. S. A. 1932, 18 (1), 1–8. https://doi.org/10.1073/pnas.18.1.1.

(56) Kang, S.-G.; Han, Z. Z.; Daude, N.; McNamara, E.; Wohlgemuth, S.; Molina-Porcel, L.; Safar, J. G.; Mok, S.-A.; Westaway, D. Pathologic Tau Conformer Ensembles Induce Dynamic, Liquid-Liquid Phase Separation Events at the Nuclear Envelope. BMC Biol. 2021, 19 (1), 199. https://doi.org/10.1186/s12915-021-01132-y.

(57) Espinoza, M.; de Silva, R.; Dickson, D. W.; Davies, P. Differential Incorporation of Tau Isoforms in Alzheimer’s Disease. J. Alzheimers Dis. JAD 2008, 14 (1), 1–16. https://doi.org/10.3233/jad-2008-14101.

(58) Anderberg, M. R. Cluster Analysis for Applications; Academic Press/Elsevier Science: Burlington, 2014.

(59) Pedregosa, F.; Varoquaux, G.; Gramfort, A.; Michel, V.; Thirion, B.; Grisel, O.; Blondel, M.; Prettenhofer, P.; Weiss, R.; Dubourg, V.; Vanderplas, J.; Passos, A.; Cournapeau, D.; Brucher, M.; Perrot, M.; Duchesnay, É. Scikit-Learn: Machine Learning in Python. J. Mach. Learn. Res. 2011, 12 (85), 2825–2830.

(60) Virtanen, P.; Gommers, R.; Oliphant, T. E.; Haberland, M.; Reddy, T.; Cournapeau, D.; Burovski, E.; Peterson, P.; Weckesser, W.; Bright, J.; van der Walt, S. J.; Brett, M.; Wilson, J.; Millman, K. J.; Mayorov, N.; Nelson, A. R. J.; Jones, E.; Kern, R.; Larson, E.; Carey, C. J.; Polat, İ.; Feng, Y.; Moore, E. W.; VanderPlas, J.; Laxalde, D.; Perktold, J.; Cimrman, R.; Henriksen, I.; Quintero, E. A.; Harris, C. R.; Archibald, A. M.; Ribeiro, A. H.; Pedregosa, F.; van Mulbregt, P.; SciPy 1.0 Contributors. SciPy 1.0: Fundamental Algorithms for Scientific Computing in Python. Nat. Methods 2020, 17 (3), 261–272. https://doi.org/10.1038/s41592-019-0686-2.

(61) Sergeant, N.; Wattez, A.; Delacourte, A. Neurofibrillary Degeneration in Progressive Supranuclear Palsy and Corticobasal Degeneration: Tau Pathologies with Exclusively “Exon 10” Isoforms. J. Neurochem. 1999, 72 (3), 1243–1249. https://doi.org/10.1046/j.1471-4159.1999.0721243.x.

(62) Umeda, Y.; Taniguchi, S.; Arima, K.; Piao, Y.-S.; Takahashi, H.; Iwatsubo, T.; Mann, D.; Hasegawa, M. Alterations in Human Tau Transcripts Correlate with Those of Neurofilament in Sporadic Tauopathies. Neurosci. Lett. 2004, 359 (3), 151–154. https://doi.org/10.1016/j.neulet.2004.01.060.

(63) Barghorn, S.; Biernat, J.; Mandelkow, E. Purification of Recombinant Tau Protein and Preparation of Alzheimer-Paired Helical Filaments in Vitro. Methods Mol. Biol. Clifton NJ 2005, 299, 35–51. https://doi.org/10.1385/1-59259-874-9:035.

(64) Aoyagi, H.; Hasegawa, M.; Tamaoka, A. Fibrillogenic Nuclei Composed of P301L Mutant Tau Induce Elongation of P301L Tau but Not Wild-Type Tau. J. Biol. Chem. 2007, 282 (28), 20309–20318. https://doi.org/10.1074/jbc.M611876200.

(65) Chen, D.; Drombosky, K. W.; Hou, Z.; Sari, L.; Kashmer, O. M.; Ryder, B. D.; Perez, V. A.; Woodard, D. R.; Lin, M. M.; Diamond, M. I.; Joachimiak, L. A. Tau Local Structure Shields an Amyloid-Forming Motif and Controls Aggregation Propensity. Nat. Commun. 2019, 10 (1), 2493. https://doi.org/10.1038/s41467-019-10355-1.

(66) Nacharaju, P.; Lewis, J.; Easson, C.; Yen, S.; Hackett, J.; Hutton, M.; Yen, S. H. Accelerated Filament Formation from Tau Protein with Specific FTDP-17 Missense Mutations. FEBS Lett. 1999, 447 (2–3), 195–199. https://doi.org/10.1016/s0014-5793(99)00294-x.

(67) Moreira, G. G.; Cristóvão, J. S.; Torres, V. M.; Carapeto, A. P.; Rodrigues, M. S.; Landrieu, I.; Cordeiro, C.; Gomes, C. M. Zinc Binding to Tau Influences Aggregation Kinetics and Oligomer Distribution. Int. J. Mol. Sci. 2019, 20 (23), 5979. https://doi.org/10.3390/ijms20235979.

(68) Zhou, Z.; Fan, J.-B.; Zhu, H.-L.; Shewmaker, F.; Yan, X.; Chen, X.; Chen, J.; Xiao, G.-F.; Guo, L.; Liang, Y. Crowded Cell-like Environment Accelerates the Nucleation Step of Amyloidogenic Protein Misfolding. J. Biol. Chem. 2009, 284 (44), 30148–30158. https://doi.org/10.1074/jbc.M109.002832.

(69) Zhong, Q.; Congdon, E. E.; Nagaraja, H. N.; Kuret, J. Tau Isoform Composition Influences Rate and Extent of Filament Formation. J. Biol. Chem. 2012, 287 (24), 20711–20719. https://doi.org/10.1074/jbc.M112.364067.

(70) Zhang, W.; Falcon, B.; Murzin, A. G.; Fan, J.; Crowther, R. A.; Goedert, M.; Scheres, S. H. Heparin-Induced Tau Filaments Are Polymorphic and Differ from Those in Alzheimer’s and Pick’s Diseases. eLife 2019, 8, e43584. https://doi.org/10.7554/eLife.43584.

(71) Chang, A.; Xiang, X.; Wang, J.; Lee, C.; Arakhamia, T.; Simjanoska, M.; Wang, C.; Carlomagno, Y.; Zhang, G.; Dhingra, S.; Thierry, M.; Perneel, J.; Heeman, B.; Forgrave, L. M.; DeTure, M.; DeMarco, M. L.; Cook, C. N.; Rademakers, R.; Dickson, D. W.; Petrucelli, L.; Stowell, M. H. B.; Mackenzie, I. R. A.; Fitzpatrick, A. W. P. Homotypic Fibrillization of TMEM106B across Diverse Neurodegenerative Diseases. Cell 2022, 185 (8), 1346–1355.e15. https://doi.org/10.1016/j.cell.2022.02.026.

(72) Sumi, S. M.; Bird, T. D.; Nochlin, D.; Raskind, M. A. Familial Presenile Dementia with Psychosis Associated with Cortical Neurofibrillary Tangles and Degeneration of the Amygdala. Neurology 1992, 42 (1), 120–127. https://doi.org/10.1212/wnl.42.1.120.

(73) Poorkaj, P.; Bird, T. D.; Wijsman, E.; Nemens, E.; Garruto, R. M.; Anderson, L.; Andreadis, A.; Wiederholt, W. C.; Raskind, M.; Schellenberg, G. D. Tau Is a Candidate Gene for Chromosome 17 Frontotemporal Dementia. Ann. Neurol. 1998, 43 (6), 815–825. https://doi.org/10.1002/ana.410430617.

(74) Rizzu, P.; Van Swieten, J. C.; Joosse, M.; Hasegawa, M.; Stevens, M.; Tibben, A.; Niermeijer, M. F.; Hillebrand, M.; Ravid, R.; Oostra, B. A.; Goedert, M.; van Duijn, C. M.; Heutink, P. High Prevalence of Mutations in the Microtubule-Associated Protein Tau in a Population Study of Frontotemporal Dementia in the Netherlands. Am. J. Hum. Genet. 1999, 64 (2), 414–421. https://doi.org/10.1086/302256.

(75) Momeni, P.; Pittman, A.; Lashley, T.; Vandrovcova, J.; Malzer, E.; Luk, C.; Hulette, C.; Lees, A.; Revesz, T.; Hardy, J.; de Silva, R. Clinical and Pathological Features of an Alzheimer’s Disease Patient with the MAPT Delta K280 Mutation. Neurobiol. Aging 2009, 30 (3), 388–393. https://doi.org/10.1016/j.neurobiolaging.2007.07.013.

(76) Goedert, M.; Spillantini, M. G.; Jakes, R.; Rutherford, D.; Crowther, R. A. Multiple Isoforms of Human Microtubule-Associated Protein Tau: Sequences and Localization in Neurofibrillary Tangles of Alzheimer’s Disease. Neuron 1989, 3 (4), 519–526. https://doi.org/10.1016/0896-6273(89)90210-9.

(77) Montgomery, K. M.; Carroll, E. C.; Thwin, A. C.; Quddus, A. Y.; Hodges, P.; Southworth, D. R.; Gestwicki, J. E. Chemical Features of Polyanions Modulate Tau Aggregation and Conformational States. J. Am. Chem. Soc. 2023, 145 (7), 3926–3936. https://doi.org/10.1021/jacs.2c08004.

(78) Lövestam, S.; Koh, F. A.; van Knippenberg, B.; Kotecha, A.; Murzin, A. G.; Goedert, M.; Scheres, S. H. W. Assembly of Recombinant Tau into Filaments Identical to Those of Alzheimer’s Disease and Chronic Traumatic Encephalopathy. eLife 2022, 11, e76494. https://doi.org/10.7554/eLife.76494.

(79) Abskharon, R.; Sawaya, M. R.; Boyer, D. R.; Cao, Q.; Nguyen, B. A.; Cascio, D.; Eisenberg, D. S. Cryo-EM Structure of RNA-Induced Tau Fibrils Reveals a Small C-Terminal Core That May Nucleate Fibril Formation. Proc. Natl. Acad. Sci. U. S. A. 2022, 119 (15), e2119952119. https://doi.org/10.1073/pnas.2119952119.

