## supplemental figures for "Disease associated mutations in tau encode for changes in aggregate structure conformation"

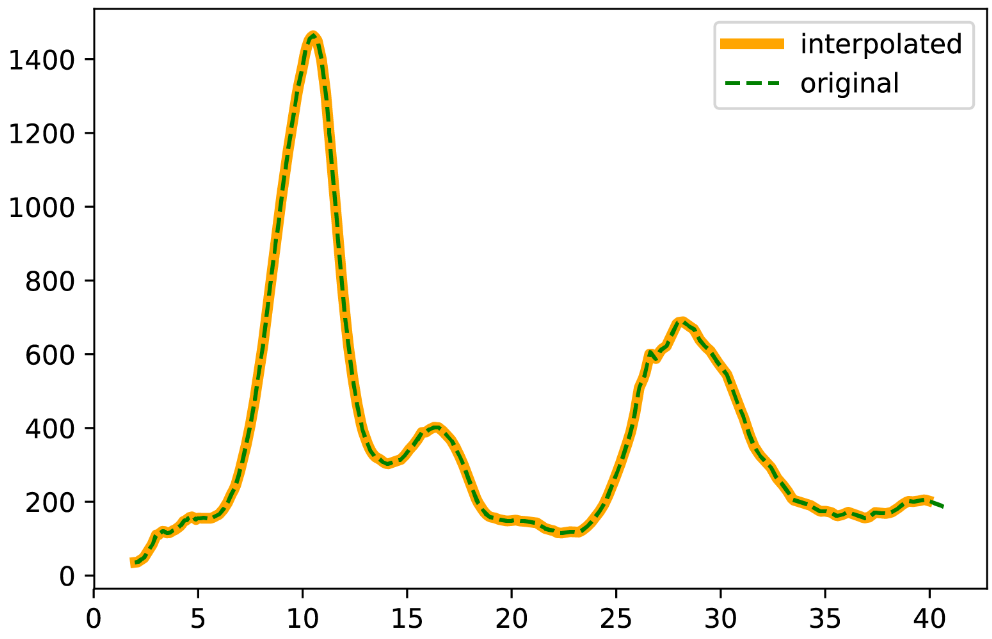

Supplementary Figure 1. **Supporting data related to analysis of trypsin-resistant tau fragment profiles.** Graph comparing the original capillary gel electrophoresis chromatogram data (green dotted line) to the data obtained after interpolation with a step size of 0.1 (yellow solid line).

**A**

plate column

| plate row | 1 | 2 | 3 | 4 | 5 |
|---|---|---|---|---|---|
| A | 1.17 | 1.08 | 1.2 | 1.05 | 0.99 |
| B | 1.07 | 0.97 | 1.2 | 1.13 | 0.89 |
| C | 0.77 | 1.01 | 1.2 | 0.92 | 0.78 |
| D | 0.7 | 0.75 | 1.01 | 0.56 | 0.57 |
| E | 0.79 | 0.78 | 0.96 | 0.86 | 0.42 |
| F | 0.84 | 0.91 | 1.14 | 1.03 | |
| G | 1.08 | 1.08 | 1.44 | 0.79 | |
| H | 0.87 | 0.84 | 1.1 | 0.79 | |

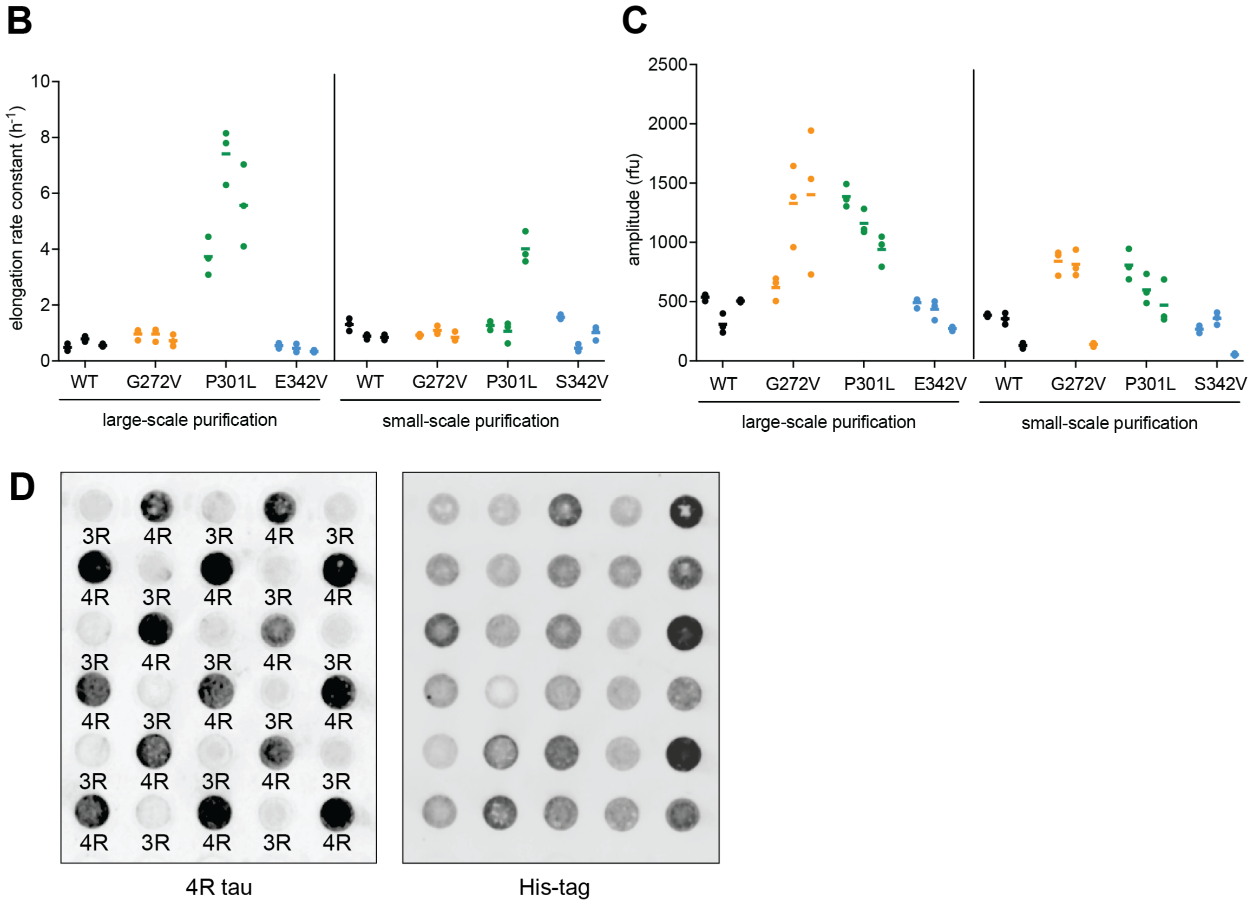

Supplementary Figure 2. **Supporting data related to Figure 1**. A) The calculated concentration in µg/µl (total volume ~ 200-400 µL) for tau proteins purified in individual wells using our small-scale purification method. Wells with values that fall within 1 STDEV of the average concentration (0.93µg/µl) or above are shaded in grey. Results representative of 3 independent experiments. B-C) Comparing aggregation kinetics of large-scale versus small-scale purified tau (WT, G272V, P301L, E342V) with respect to B) elongation rate constant and C) amplitude. Individual data points for three technical replicates (aligned dots), assayed from three individual protein preparations (side-by-side columns of dots) are plotted along with the mean ± SD for each experiment (bar). D) Replicate dot blots of small-scale purified 0N4R and 0N3R tau probed with a 4R tau antibody (left image) or a His-tag antibody (right image). Each dot is loaded with a purified tau sample. The relative location of the purified tau sample in the dot blot and purification plate is preserved. The identity of the tau sample being purified in each well (0N4R or 0N3R) is indicated (4R or 3R).

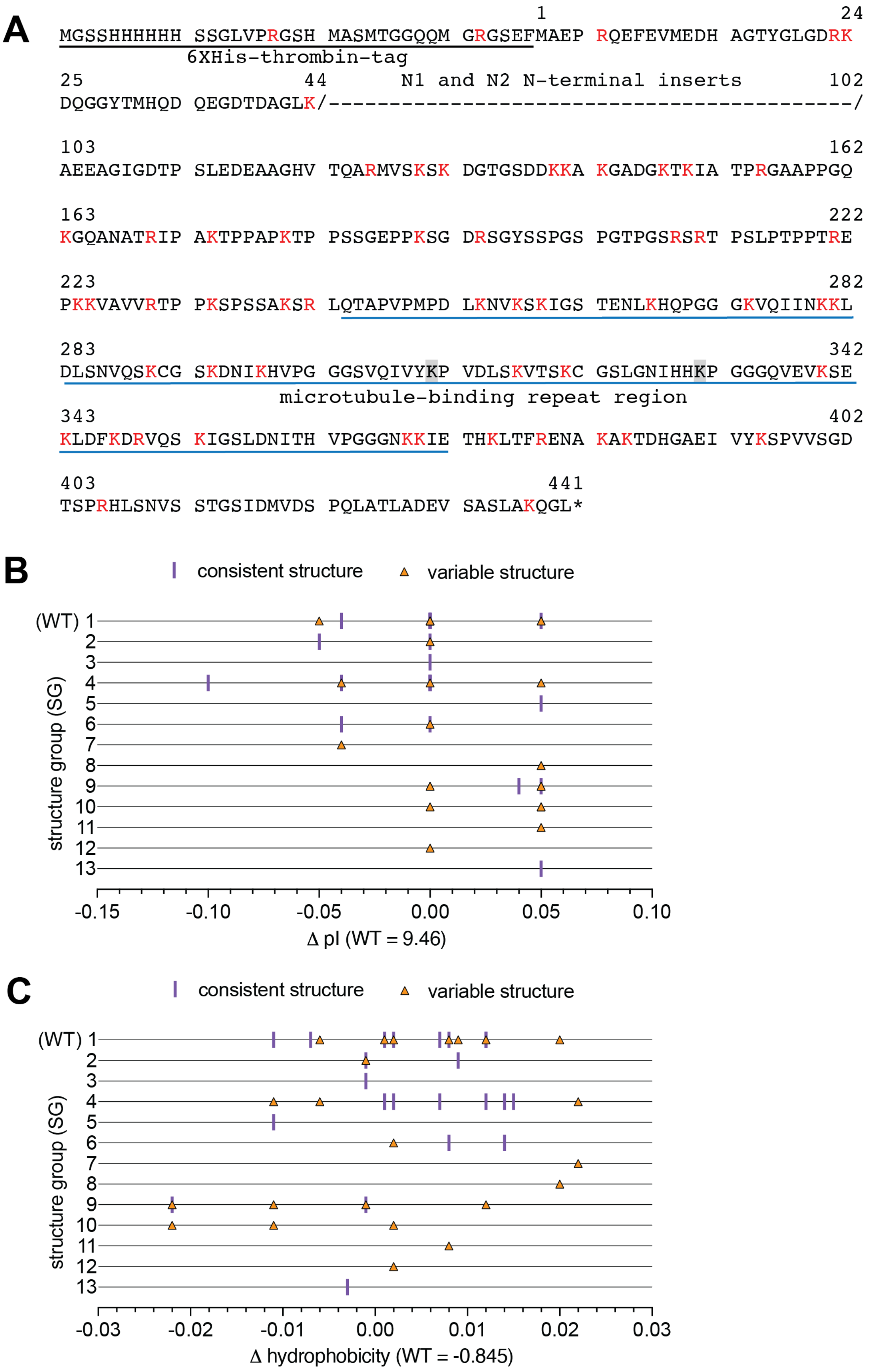
A
1
24
MGSSHHHHHH SSGLVPRGSH MASMTGGQQM GRGSEFMAEP RQEFEVMEDH AGTYGLGDRK
6XHis-thrombin-tag
25
44
N1 and N2 N-terminal inserts
102
DQGGYTMHQD QEGDTDAGLK/------------------------------------------/
103
162
AEEAGIGDTP SLEDEAAGHV TQARMVSKSK DGTGSDDKKA KGADGKTKIA TPRGAAPPGQ
163
222
KGQANATRIP AKTPPAPKTP PSSGEPPKSG DRSGYSSPGS PGTPGSRSRT PSLPTPPTRE
223
282
PKKVAVVRTP PKSPSSAKSR LQTAPVPMPD LKNVKSKIGS TENLKHQPGG GKVQIINKKL
283
342
DLSNVQSKCG SKDNIKHVPG GGSVQIVYKP VDLSKVTSKC GSLGNIHHKP GGGQVEVKSE
microtubule-binding repeat region
343
402
KLDFKDRVQS KIGSLDNITH VPGGGNKKIE THKLTFRENA KAKTDHGAEI VYKSPVVSGD
403
441
TSPRHLSNVS STGSIDMVDS PQLATLADEV SASLAKQGL*
B
consistent structure
variable structure
(WT) 1
structure group (SG)
-0.15
-0.10
-0.05
0.00
0.05
0.10
Δ pI (WT = 9.46)
C
consistent structure
variable structure
(WT) 1
structure group (SG)
-0.03
-0.02
-0.01
0.00
0.01
0.02
0.03
Δ hydrophobicity (WT = -0.845)

Supplementary Figure 3. **Supporting data related to Figure 2.** A) Amino acid sequence of N-terminal His-thrombin tagged WT 0N4R tau. Numbering of the tau sequence is based on the conventional 2N4R numbering system (1-441). The location of missing N1 and N2 N-terminal insert sequences caused during splicing of 0N4R transcript is indicated by a dashed line region (/ - - - /). Trypsin cleavage sites (K/R) are colored in red. Trypsin sites that are blocked due to specific P2, P1' identities are highlighted in grey. The sequence corresponding to His-thrombin tag is underlined in black. The sequence region corresponding to the microtubule binding repeat region (R1-R4) is underlined in blue. B - C) 0N4R tau mutants plotted according to their structure group classification (y-axis) and on the x-axis their change from WT with respect to B) isoelectric point or C) hydrophobicity. Tau mutants that gave a single digest profile across three trials were termed as having a consistent structure (purple line) whereas those that gave an alternate profile in at least one experiment were classified as having variable structure (orange triangles, all structure groups indicated for each mutant).

A

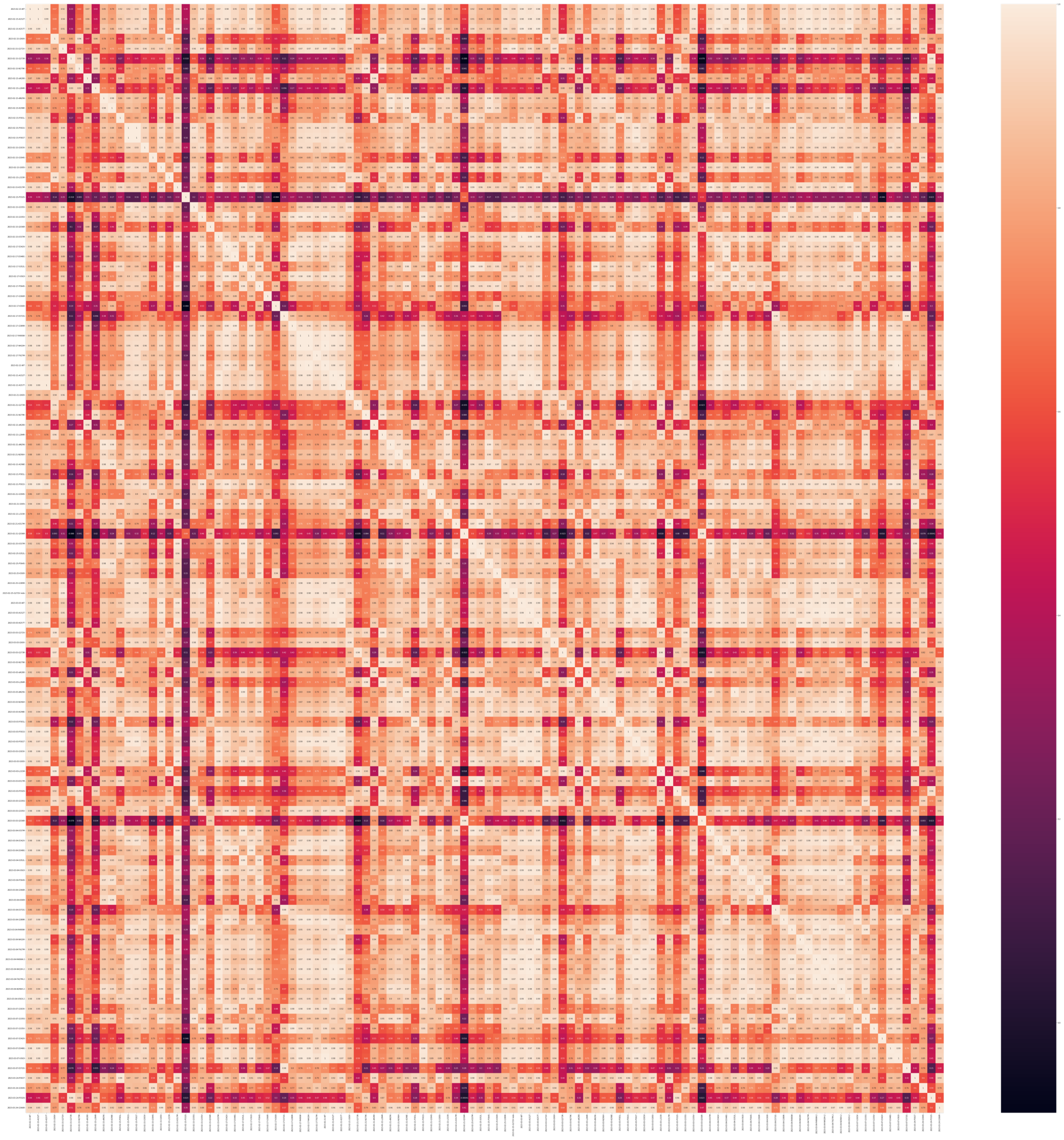

**B**

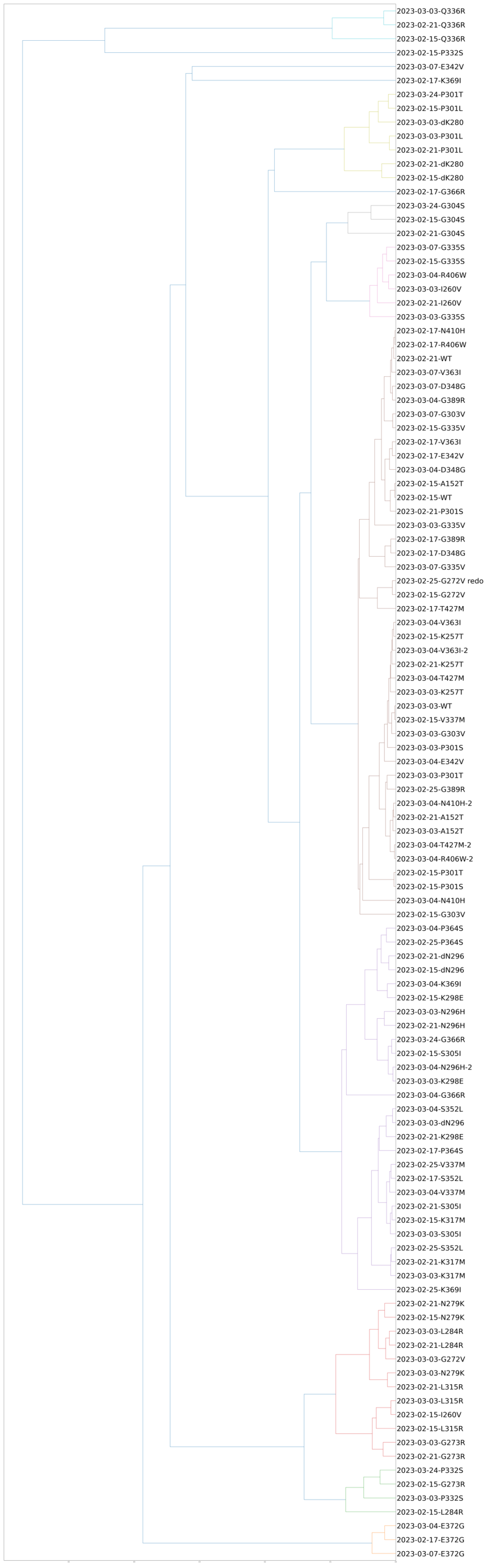
2023-03-03-Q336R
2023-02-21-Q336R
2023-02-15-Q336R
2023-02-15-P332S
2023-03-07-E342V
2023-02-17-K369I
2023-03-24-P301T
2023-02-15-P301L
2023-03-03-dK280
2023-03-03-P301L
2023-02-21-P301L
2023-02-21-dK280
2023-02-15-dK280
2023-02-17-G366R
2023-03-24-G304S
2023-02-15-G304S
2023-02-21-G304S
2023-03-07-G335S
2023-02-15-G335S
2023-03-04-R406W
2023-03-03-I260V
2023-02-21-I260V
2023-03-03-G335S
2023-02-17-N410H
2023-02-17-R406W
2023-02-21-WT
2023-03-07-V363I
2023-03-07-D348G
2023-03-04-G389R
2023-03-07-G303V
2023-02-15-G335V
2023-02-17-V363I
2023-02-17-E342V
2023-03-04-D348G
2023-02-15-A152T
2023-02-15-WT
2023-02-21-P301S
2023-03-03-G335V
2023-02-17-G389R
2023-02-17-D348G
2023-03-07-G335V
2023-02-25-G272V redo
2023-02-15-G272V
2023-02-17-T427M
2023-03-04-V363I
2023-02-15-K257T
2023-03-04-V363I-2
2023-02-21-K257T
2023-03-04-T427M
2023-03-03-K257T
2023-03-03-WT
2023-02-15-V337M
2023-03-03-G303V
2023-03-03-P301S
2023-03-04-E342V
2023-03-03-P301T
2023-02-25-G389R
2023-03-04-N410H-2
2023-02-21-A152T
2023-03-03-A152T
2023-03-04-T427M-2
2023-03-04-R406W-2
2023-02-15-P301T
2023-02-15-P301S
2023-03-04-N410H
2023-02-15-G303V
2023-03-04-P364S
2023-02-25-P364S
2023-02-21-dN296
2023-02-15-dN296
2023-03-04-K369I
2023-02-15-K298E
2023-03-03-N296H
2023-02-21-N296H
2023-03-24-G366R
2023-02-15-S305I
2023-03-04-N296H-2
2023-03-03-K298E
2023-03-04-G366R
2023-03-04-S352L
2023-03-03-dN296
2023-02-21-K298E
2023-02-17-P364S
2023-02-25-V337M
2023-02-17-S352L
2023-03-04-V337M
2023-02-21-S305I
2023-02-15-K317M
2023-03-03-S305I
2023-02-25-S352L
2023-02-21-K317M
2023-03-03-K317M
2023-02-25-K369I
2023-02-21-N279K
2023-02-15-N279K
2023-03-03-L284R
2023-02-21-L284R
2023-03-03-G272V
2023-03-03-N279K
2023-02-21-L315R
2023-03-03-L315R
2023-02-15-I260V
2023-02-15-L315R
2023-03-03-G273R
2023-02-21-G273R
2023-03-24-P332S
2023-02-15-G273R
2023-03-03-P332S
2023-02-15-L284R
2023-03-04-E372G
2023-02-17-E372G
2023-03-07-E372G

Supplementary Figure 4. **Supporting data related to Figure 2 - Binning of structure groups**. Comparative analysis of digest fragment profiles across all samples (WT and mutant tau variants, 3 samples per variant) generates the outputs: A) heatmap representing the computed distance between individual samples and B) dendrogram that groups samples by hierarchical clustering using average linkage. A 90% similarity threshold was used to bin samples into 13 structure groups as described in Fig. 2 and Table 2.

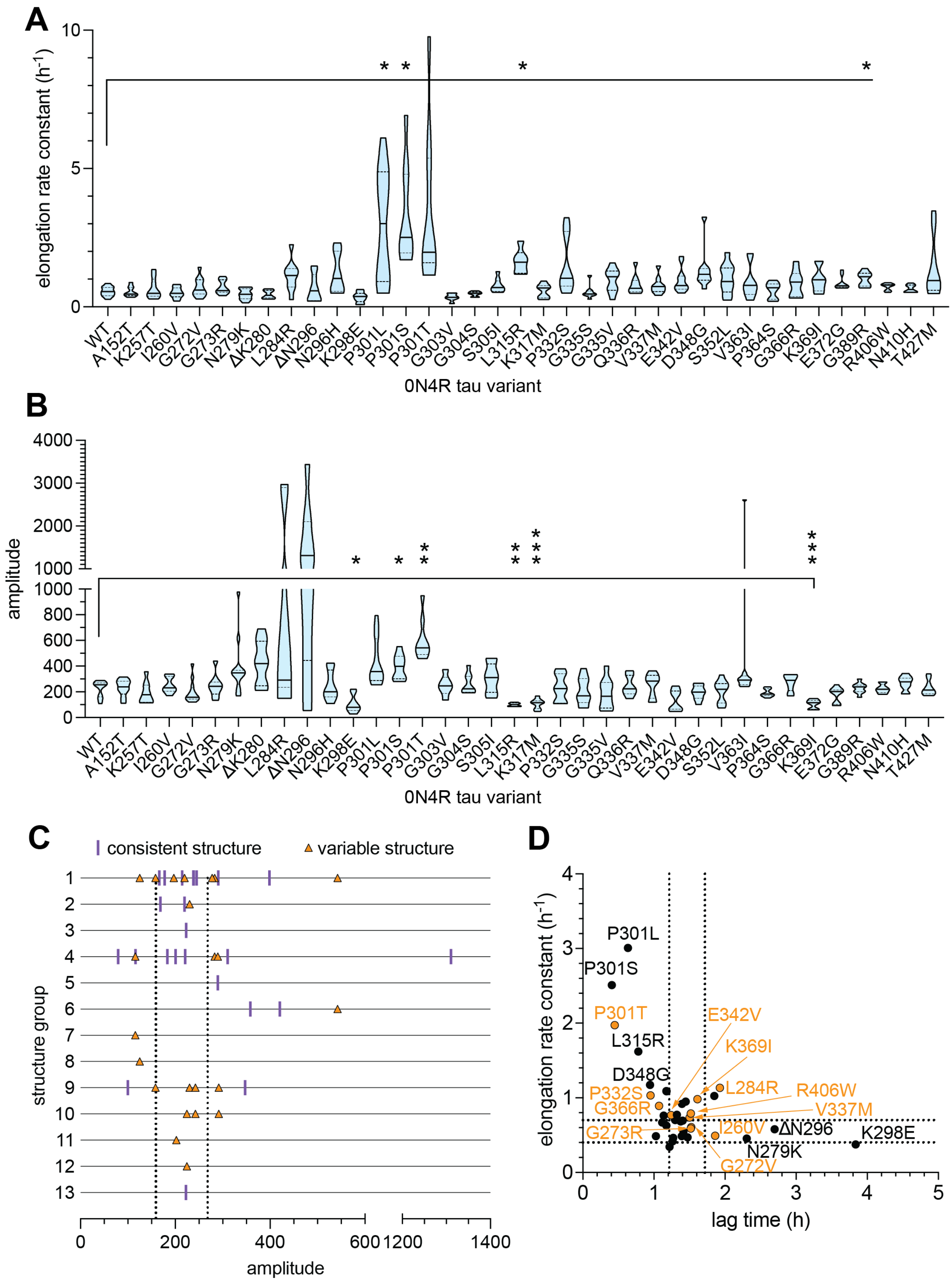

A
elongation rate constant (h-1)
0N4R tau variant
WT A152T K257T I260V G272V G273R N279K ΔK280 L284R ΔN296 N296H K298E P301L P301S P301T G303V G304S S305I L315R K317M P332S G335S G335V Q336R V337M E342V D348G S352L V363I P364S G366R K369I E372G G389R R406W N410H T427M
B
amplitude
0N4R tau variant
C
consistent structure
variable structure
structure group
amplitude
D
elongation rate constant (h-1)
lag time (h)
P301L
P301S
P301T
L315R
D348G
P332S
G366R
G273R
E342V
K369I
L284R
R406W
V337M
I260V
ΔN296
K298E
N279K
G272V

Supplementary Figure 5. **Supporting data related to Figure 3.** Fitted values from aggregation kinetic curves for A) elongation rate constant and B) amplitude of individual tau WT and mutants. The median (solid line) and quartiles (dotted lines) are plotted for each variant ($n \geq 9$ replicates, $\geq 3$ independent experiments). A Brown-Forsythe and Welch ANOVA analysis was performed with post-hoc comparisons to the WT tau group. p =* <0.05, **<0.01, ***<0.001. C) 0N4R tau mutants plotted according to their calculated amplitude values (x-axis, median, $n \geq 9$ replicates, $\geq 3$ independent experiments) and structure group classification (y-axis).Tau mutants that gave a single digest profile across three trials were termed as having a consistent structure (purple line) whereas those that gave an alternate profile in at least one experiment were classified as having variable structure (orange triangles, all structure groups indicated for each mutant). D) Comparison of all tau variants with respect to: elongation rate constant, lag time , and consistent or variable structure group classification. For each tau variant, median lag time and elongation rate constant values are plotted ($n \geq 9$ replicates, $\geq 3$ independent experiments). Tau mutants that gave a single digest profile across three trials were termed as having a consistent structure (black circles) whereas those that gave an alternate profile in at least one experiment were classified as having variable structure (orange circles). For WT tau, the quartile boundaries of lag time and elongation rate constant values are marked by dashed lines.

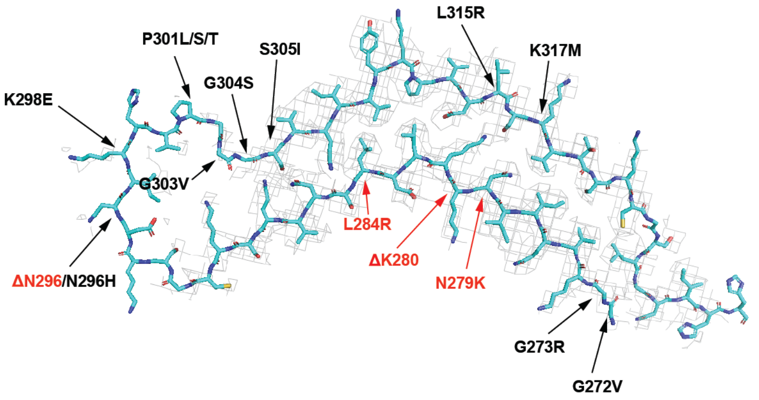

Supplementary Figure 6. **Cryo-EM density map (grey mesh) of snake filament derived from recombinant 2N4R tau aggregation with heparin (PDB: 6QJH)**. The backbone trace is colored in blue. Disease associated mutations assayed in our study are labeled according to whether they are predicted to cause a steric clash (red) or not (black) as indicated in Fig. 4A and Table 2.

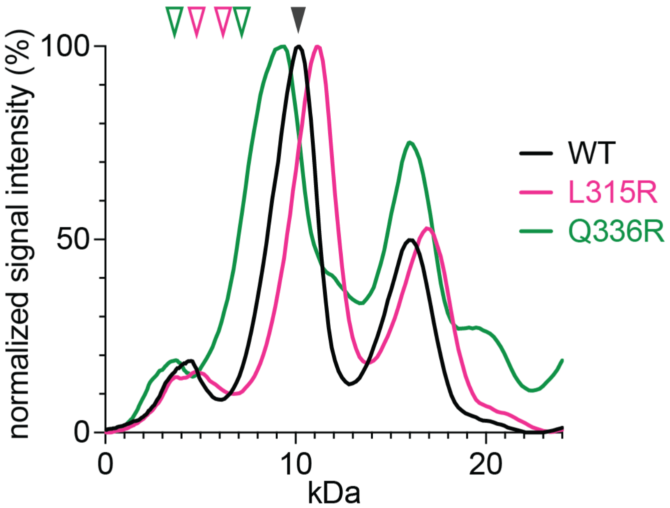

Supplementary Figure 7. **Altered digest profiles of tau aggregate mutants is not due to addition of tryptic sites by K or R mutations**. Representative chromatograms of digested fragment profiles of WT, L315R and Q336R tau mutants over the 0-24 kDa region. For the y-axis, signal intensity is normalized with the maximal signal value set to 100%. The black arrowhead indicates the 10kDa fragment from the WT-tau digest profile (apparent MW=10 kDa, calculated MW=11kDa) which has been mapped to cover sequence regions from approximately residues 267-375. If L315R and Q336R are proposed to retain a WT-tau aggregate structure, any altered cleavage at the mutated residues would result in fragmentation of the 10 kDa band. Apparent MW of fragments created by cleavage at mutated residues are indicated with colored open arrowheads (calculated fragment MW for L315R= 6.1 and 4.7 kDa, calculated fragment MW for Q336R=7.2 and 3.5 kDa).
